# *Polarella glacialis* genomes encode tandem repeats of single-exon genes with functions critical to adaptation of dinoflagellates

**DOI:** 10.1101/704437

**Authors:** Timothy G. Stephens, Raúl A. González-Pech, Yuanyuan Cheng, Amin R. Mohamed, David W. Burt, Debashish Bhattacharya, Mark A. Ragan, Cheong Xin Chan

**Affiliations:** Institute for Molecular Bioscience, The University of Queensland, Brisbane, QLD 4072, Australia; UQ Genomics Initiative, The University of Queensland, Brisbane, QLD 4072, Australia; Commonwealth Scientific and Industrial Research Organisation (CSIRO) Agriculture and Food, Queensland Bioscience Precinct, Brisbane, QLD 4067, Australia; UQ Genomics, The University of Queensland, Brisbane, QLD 4072, Australia; Department of Biochemistry and Microbiology, Rutgers University, New Brunswick, NJ 08901, U.S.A.; School of Chemistry and Molecular Biosciences, The University of Queensland, Brisbane, QLD 4072, Australia

## Abstract

Dinoflagellates are taxonomically diverse, ecologically important phytoplankton in marine and freshwater environments. Here, we present two draft diploid genome assemblies of the free-living dinoflagellate *Polarella glacialis*, isolated from the Arctic and Antarctica. For each genome, guided using full-length transcriptome data, we predicted >50,000 high-quality genes. About 68% of the genome is repetitive sequence; long terminal repeats likely contribute to intra-species structural divergence and distinct genome sizes (3.0 and 2.7 Gbp). Of all genes, ∼40% are encoded unidirectionally, ∼25% comprised of single exons. Multi-genome comparison unveiled genes specific to *P. glacialis* and a common, putatively bacterial, origin of ice-binding domains in cold-adapted dinoflagellates. Our results elucidate how selection acts within the context of a complex genome structure to facilitate local adaptation. Since most dinoflagellate genes are constitutively expressed, *Polarella glacialis* has enhanced transcriptional responses *via* unidirectional, tandem duplication of single-exon genes that encode functions critical to survival in cold, low-light environments.

## Introduction

Dinoflagellates are a species-rich (2,270 species in AlgaeBase^1^) and anciently diverged (likely Precambrian origin^2^) group of phytoplankton that are ubiquitous in marine and fresh waters. Mostly photosynthetic, dinoflagellates form the base of food webs. They sustain global aquatic ecosystems *via* primary production and cycling of organic carbon and nitrogen. Some dinoflagellate lineages comprise species that are symbiotic or parasitic. For example, members of the family Symbiodiniaceae are crucial symbionts in corals and other coral reef animals^3, 4^, and parasitic dinoflagellates can cause death in economically important crustaceans, such as crabs and lobsters^5^. Most dinoflagellates, however, are free-living.

Bloom-forming taxa may cause “red tides”, which produce toxins that pose serious human health risks^6^. Some taxa have specialised to inhabit extreme environments, such as those found in the brine channels of polar sea ice^7–10^.

Thus far, available genome data of dinoflagellates are largely restricted to symbiotic or parasitic species^11–17^. These lineages were chosen for sequencing because their genomes are relatively small, i.e. 0.12-4.8 Gbp. In comparison, genomes of other free-living dinoflagellates are much larger in size, ranging from ∼7 Gbp in the psychrophile *Polarella glacialis*, to over 200 Gbp in *Prorocentrum* sp. based on DAPI-staining of DNA content^18^.

Repeat content has been estimated at >55% in the genome sequences of some free-living dinoflagellates^19, 20^; single-exon genes have also been described^21^. Given that most dinoflagellate lineages are free-living, whole genome sequences of these taxa are critical to understand the molecular mechanisms that underpin their successful diversification in specialised environmental niches.

*Polarella glacialis*, a psychrophilic (cold-adapted) free-living species, represents an excellent system for genomic studies of dinoflagellates for three reasons. First, it is closely related to Symbiodiniaceae (both in Order Suessiales), the family that contains the coral reef symbionts, *e.g. Symbiodinium* and related genera. Second, *P. glacialis* has been reported only in polar regions. Studying the *P. glacialis* genome can thus provide a first glimpse into molecular mechanisms that underlie both the evolutionary transition of dinoflagellates from a free-living to a symbiotic lifestyle, and the adaptation to extreme environments. Third, the estimated genome size of *P. glacialis* is still in the smaller range (∼7 Gbp^18^) of all dinoflagellate taxa, which presents a technical advantage in terms of allowing efficient genome assembly and gene prediction.

Here, we report draft *de novo* genome sequences from two *P. glacialis* isolates: CCMP1383 and CCMP2088. The former is a xenic culture first isolated from brine in the upper sea ice in McMurdo Sound (Ross Sea, Antarctica) in 1991^10^, and the latter is a xenic culture first isolated from a water sample collected adjacent to ice in northern Baffin Bay in 1998^22^.

These genomes represent the first generated from any free-living, psychrophilic dinoflagellates. Incorporating full-length transcriptome data, we investigated gene structure, repeat content, and intra-species genome divergence. Our results reveal remarkable difference in genome sizes between these two isolates of the same species, and provide evidence of tandemly repeated, single-exon genes in shaping the evolution of dinoflagellate genomes.

## Results

### Genomes of *Polarella glacialis*

Draft genome assemblies for two *Polarella glacialis* isolates (CCMP1383 and CCMP2088) were generated using a combination of Illumina short-read and PacBio long-read data (Table 1 and Supplementary Table 1). Both genomes appear diploid based on their bimodal distributions of *k*-mer counts observed from the sequence data, which closely (model fit >92%) matches the standard theoretical diploid model (Fig. 1a and Supplementary Fig. 1). These are the first diploid genomes reported for any dinoflagellate. As no reliable haploid representation could be generated (see Methods), we used these diploid assemblies in subsequent analyses. The CCMP1383 assembly had fewer and more-contiguous scaffolds (33,494; N50 length 170 Kbp; Table 1) compared to the CCMP2088 assembly (37,768; N50 length 129 Kbp; Table 1); this is likely a consequence of more long-read data generated for CCMP1383 (Supplementary Table 2). Both assemblies are much more contiguous than their corresponding assemblies generated using only short-read data (N50 length < 73 Kbp; Supplementary Table 1). For CCMP1383 and CCMP2088 respectively, the total diploid assembly sizes are 2.98 Gbp and 2.76 Gbp (Table 1 and Supplementary Table 1), and are very similar to independent diploid genome-size estimates of 3.02 Gbp and 2.65 Gbp (Supplementary Table 3). These genomes are smaller than previously estimated (∼7 Gbp based on staining of total [assumed haploid, but potentially diploid] DNA content^18^).

**Fig. 1.**
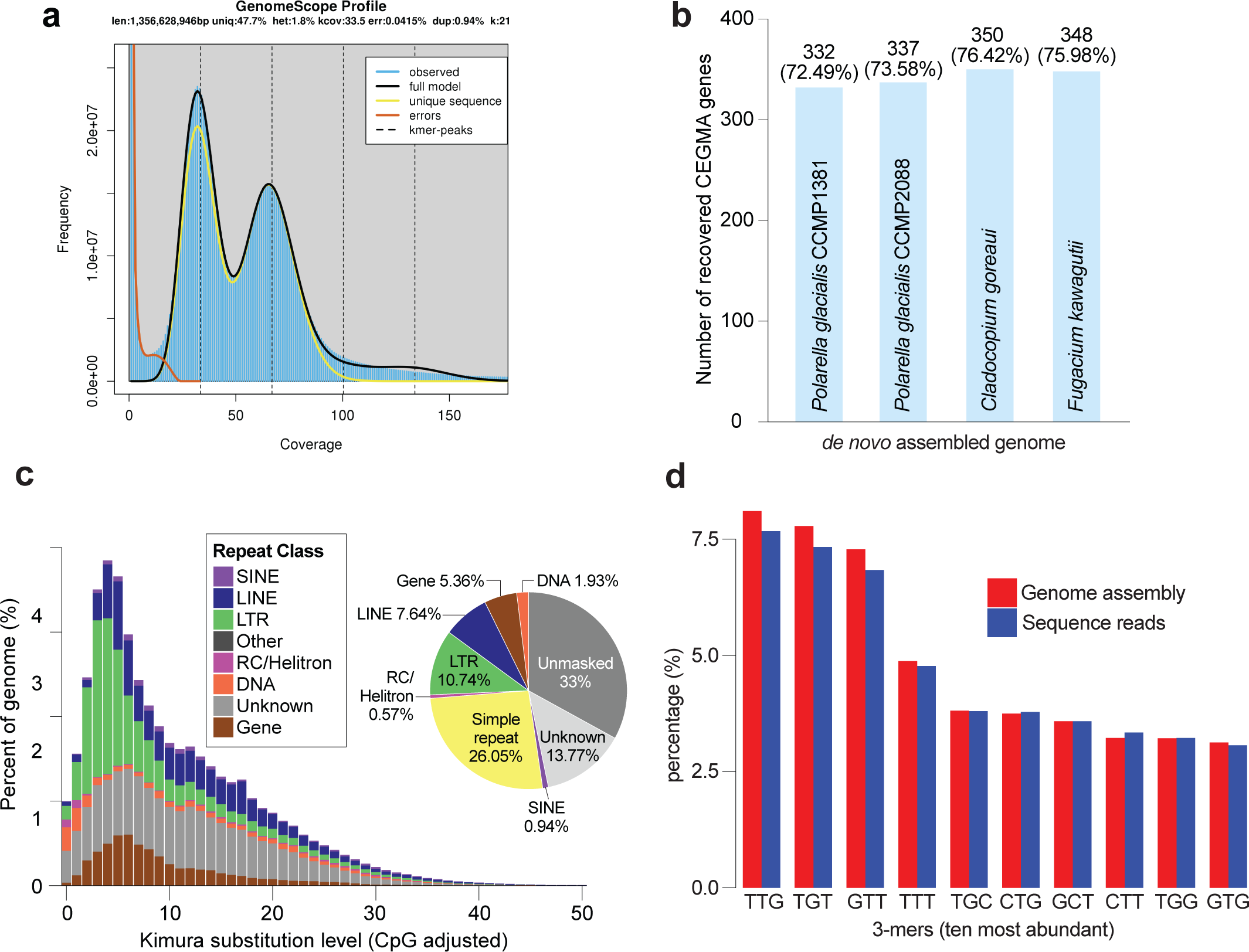
Genomes of *Polarella glacialis* and repeat content. (a) GenomeScope 21-mer profile for CCMP1383. (b) Identification of conserved core eukaryote genes (using CEGMA) in the assembled *P. glacialis* genomes of CCMP1383 and CCMP2088 compared to the assembled genomes of *Cladocopium goreaui* and *Fugacium kawagutii*^12^. (c) Interspersed repeat landscape and proportion of distinct repeat classes in the assembled genome of CCMP1383, studied using sequence divergence under the Kimura evolutionary model. (d) Percentage of 3-mers in the assembled genome and the sequence data for CCMP1383 for the ten most-abundant 3-mers.

**Table 1.** Assembled genomes of P. glacialis compared to representative publicly available dinoflagellate genomes. A more-comprehensive summary including all other available genomes is shown in Supplementary Table 1. Estimated diploid genome size for *P. glacialis* isolates shown in brackets.

|  | *Polarella glacialis* |  | Symbiodiniaceae |  |  |  | Parasitic |  |
| --- | --- | --- | --- | --- | --- | --- | --- | --- |
|  | CCMP1383 | CCMP2088 | *Symbiodinium microadriaticum* | *Breviolum minutum* | *Cladocopium goreau* | *Fugacium kawagutii* | *Amoebophrya ceratii* | *Hematodinium* sp. |
| Reference | This study | This study | Aranda et al.<sup>13</sup> | Shoguchi et al.<sup>14</sup> | Liu et al.<sup>12</sup> | Liu et al.<sup>12</sup> | John et al.<sup>16</sup> | Gornik et al.<sup>17</sup> |
| %G+C | 45.91 | 46.15 | 50.51 | 43.46 | 44.83 | 45.72 | 55.92 | 47.31 |
| Total number of scaffolds | 33,494 | 37,768 | 9,695 | 21,899 | 41,289 | 16,959 | 2,351 | 869,500 |
| Total assembled bases (Gbp) | 2.98 | 2.76 | 0.81 | 0.61 | 1.03 | 1.05 | 0.09 | 4.77 |
| N50 length of scaffolds (bp) | 170,304 | 129,205 | 573,512 | 125,226 | 98,034 | 268,823 | 83,970 | 17,235 |
| Maximum scaffold length (bp) | 2,170,995 | 1,500,384 | 3,144,590 | 810,747 | 8,337,000 | 5,159,000 | 536,776 | 186,000 |
| Estimated genome size (Gbp) | 1.48 (3.02) | 1.30 (2.65) | 1.10 | 1.5 | 1.19 | 1.07 | 0.12 | 4.8 |

However, they remain larger than those of Symbiodiniaceae^11–15^ (haploid; between 1.1 and 1.5 Gbp), and smaller than the 4.8 Gbp genome of the parasitic *Hematodinium* sp.^17^ (Table 1 and Supplementary Table 1). Our results reaffirm the tendency of DNA staining or flow cytometry to overestimate genome sizes of dinoflagellates^11–15^, potentially due to the liquid crystal structure of dinoflagellate chromosomes^23^.

The non-repetitive regions from both assembled genomes are almost identical in content and sequence. In a comparison between the non-repetitive regions of CCMP2088 against the genome of CCMP1388, top hits cover 96.9% of the query bases with an average sequence identity of 99.4%. Likewise, in CCMP1383, top hits cover 95.6% of query bases with an average sequence identity of 99.2% compared against the CCMP2088 genome. Remarkably, the estimated diploid genome size of the Antarctic isolate (CCMP1383) is approximately 370 Mbp larger than that of the Arctic isolate (CCMP2088). These results reveal, for the first time, structural divergence of genomes in dinoflagellates even within a single species, potentially explained by the uneven expansion of repetitive elements (see below). The two genome assemblies are reasonably complete. Similar proportions of core conserved eukaryote genes were recovered, i.e. 332 (72.49%) and 337 (73.58%) of the 458 CEGMA^24^ genes in CCMP1383 and CCMP2088, respectively (Fig. 1b and Supplementary Table 4). These numbers are comparable to those recovered in published symbiodiniacean genomes^12^, e.g. 350 in *C. goreaui* and 348 in *F. kawagutii*, analysed using the same approach (Fig. 1b).

The high extent of genome-sequence similarity between the two geographically distinct *P. glacialis* isolates suggests that they have either recently been transferred from one polar region to the other, or are being actively transported between the two locations, allowing for mixing of the two populations. To identify if *P. glacialis* is being actively transported between polar regions we interrogated the TARA Oceans database for the presence of this species in the broadly sampled list of sites. Despite *P. glacialis* sequences being listed in 67 of the 68 TARA sample locations, an exhaustive search based on sequence similarity did not recover clear evidence of any *P. glacialis* sequences (see Methods). Whereas we cannot dismiss the presence of *P. glacialis* at low, undetectable levels, we find no evidence at this time to support the presence of *P. glacialis* in the waters outside of the polar regions.

### *Polarella glacialis* genomes are highly repetitive

Both *P. glacialis* genomes reveal a high content of repetitive elements that encompass ∼68% (by length) of the assembled sequences (Fig. 1c and Supplementary Fig. 2). Most of these elements are simple and unknown repeats (i.e., unclassified *de novo* repeats, covering ∼13.5% of each assembled genome; Fig. 1c). The proportion of repeats in *P. glacialis* genomes is more than two-fold higher than that reported in Symbiodiniaceae (e.g., 27.9% in *Symbiodinium microadriaticum*, 16% in *Fugacium kawagutii*)^12^. This observation is not unexpected, because even before high-throughput sequencing technology was available, the genome of the biotechnologically important dinoflagellate *Crypthecodinium cohnii* was estimated to contain 55-60% repeat content^19^. A genome survey of *Alexandrium ostenfeldii* estimated the repeat content at ∼58%^20^. In comparison, the genome surveys of *Heterocapsa triquetra*^25^ and *Prorocentrum minimum*^26^ estimated their repeat content at only ∼5% and ∼6%, respectively. These values are likely underestimates because only 0.0014% of the *H. triquetra* genome was surveyed, and >28% of the reported *P. minimum* genome data is putatively of bacterial origin^26^.

The prevalence of repeats in *P. glacialis* genomes may explain their larger genome sizes compared to symbiotic dinoflagellates^11–15^, and may represent a genome signature of free-living dinoflagellates. These repeats are more conserved in *P. glacialis* (Kimura substitution level^27^ centred around 5; Fig. 1c and Supplementary Fig. 2) than those reported in Symbiodiniaceae (Kimura substitution level 10-30^12^). We also recovered a substantial proportion of long-terminal repeat (LTR) elements (∼12%) in *P. glacialis* genomes; these elements were largely absent (<0.7%) in Symbiodiniaceae^12^. Transposable elements (such as LTRs) commonly comprise up to 80% of the genomes of plants and are induced by genome shock and polyploidization, resulting in genome restructuring^28^. The abundance of LTRs in *P. glacialis* and the role of LTRs in genome restructuring may explain in part the difference in genome sizes between the two isolates. These results suggest that repetitive elements and LTRs are key contributors that drive genome size evolution of *P. glacialis*, both as a free-living and a cold-adapted dinoflagellate species. Because available dinoflagellate genomes (e.g., of Symbiodiniaceae) thus far have been generated largely using Illumina short-read data, we cannot dismiss the possibility that mis-assembly (an inevitable artefact with short-read data) may have caused the under-estimation of repeat content (and the apparent absence of LTRs) in these genomes.

In an independent analysis of simple repeats (see Methods), 25.01% and 24.17% of the CCMP1383 and CCMP2088 genomes, respectively, are found to be composed of simple repeats. The most prominent simple repeat is the trinucleotide (TTG)n (in all six reading frames; see Methods) that covered 19.1% and 18.5% of the CCMP1383 and CCMP2088 genome assemblies, respectively. The proportion of (TTG)n, observed as possible 3-mers of TTG, TGT or GTT (each ∼7-8%) in the assembled genomes, is very similar to that observed in the sequence-read data (Fig. 1d and Supplementary Table 5). Therefore, this observed prevalence of (TTG)n is unlikely due to assembly artefacts.

### DinoSL in full-length transcripts of *Polarella glacialis*

To generate high-quality supporting data to guide our gene-prediction workflow, we generated transcriptomes from both *P. glacialis* isolates, including full-length transcripts using PacBio IsoSeq technology (see Methods). Mature nuclear transcripts of dinoflagellates are known to contain a 22-nucleotide trans-spliced leader sequence (DinoSL: DCCGTAGCCATTTTGGCTCAAG, where D = T, A, or G) at the 5′-end^29^. Relic DinoSL sequences arise when transcripts with attached DinoSL are integrated back into the genome, expressed and trans-spliced with a new leader sequence^30^. Successive rounds of transcript re-integration result in multiple relic DinoSLs on a single transcript. We searched the full-length transcripts (435,032 in CCMP1383 and 1,266,042 in CCMP2088) for the presence of DinoSL and relic DinoSL sequences (see Methods). DinoSL sequences were recovered in 13.54% and 50.39% of transcripts (hereinafter DinoSL-type transcripts) in CCMP1383 and CCMP2088, respectively (Supplementary Table 6). An earlier study^31^ reported a single transcript in CCMP2088 that has a non-canonical DinoSL sequence (ATCGTAGCCATGTTGGCTCAAG), but our exhaustive search against the transcripts in either isolate did not recover this sequence.

Although our experiment (see Methods) was designed to recover full-length transcripts (with complete 5′ and 3′ regions), it is possible that the adopted library preparation step was suboptimal and the lack of a DinoSL in many transcripts may be explained by mRNA degradation. This may, in the first instance, be reflected by the varying degrees of truncation of the DinoSL-type transcripts in our data. However, 70.45% of these transcripts in CCMP1383 and 69.18% of transcripts in CCMP2088 start at one of the two contiguous cytosine bases (i.e. at positions 2 and 8) of the DinoSL (Fig. 2a; Supplementary Table 7).

**Fig. 2.**
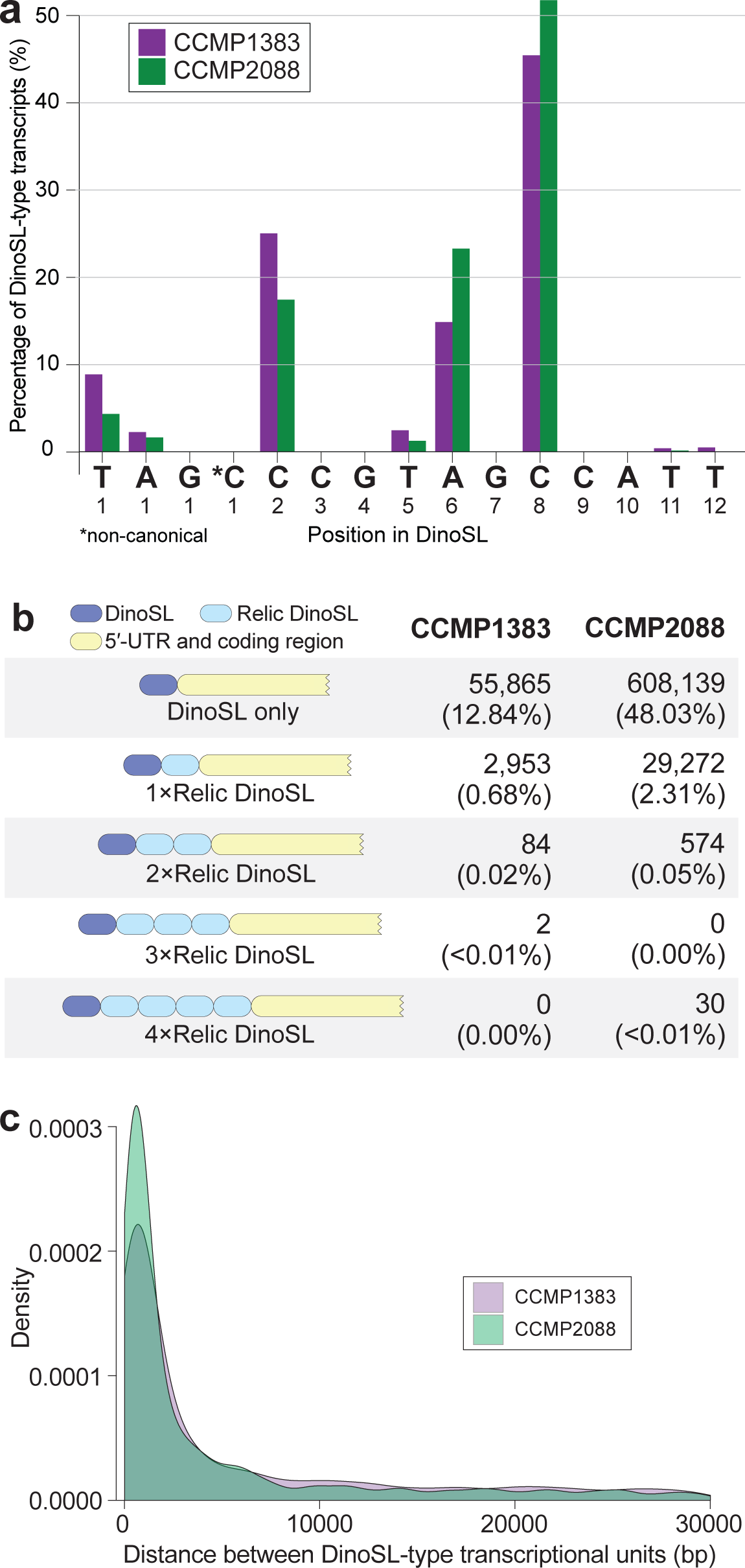
DinoSL-type full-length transcripts in *P. glacialis*. (a) Percentage of DinoSL-type transcripts of *P. glacialis* based on the identified start position along the DinoSL sequence, shown for positions 1 through 12. (b) Structure and number of DinoSL and/or relic DinoSL containing IsoSeq transcripts from each isolate. (c) Distribution of distances (in bp) between DinoSL-type transcriptional units shown for transcriptomes of CCMP1383 and CCMP2088.

This preference for start sites at the double-cytosine positions suggests that the 5′ selection method we used (that purifies for the 5′ methylated cap site) is binding to these regions instead of the true 5′-cap. This in turn may happen because cytosines at these sites are methylated. Cytosine methylation has been described in genomes of eukaryotes including dinoflagellates, potentially as a mechanism for silencing of transposable elements and regulation of gene expression^32–34^. A recent study of the *Breviolum minutum* genome revealed that cytosine methylation often occurred at CG dinucleotides^35^. The impact of methylation on recovery of splice leaders in dinoflagellates remains to be systematically investigated.

In CCMP1383 and CCMP2088, 0.68% and 2.31% of all full-length transcripts respectively were found to encode one relic DinoSL (immediately following their primary DinoSL) while smaller proportions (0.020% and 0.048% respectively) encode multiple relic DinoSL sequences (Fig. 2b). We recovered 30 transcripts in CCMP2088 that have four putative relic DinoSL sequences; they shared >99% sequence identity among one another. Five of these 30 transcripts are shorter than the others, suggesting an alternative transcription 3′-termination site, thus a distinct isoform.

We further assessed the diversity of alternative splice-forms by clustering the full-length transcripts by sequence similarity using PASA (see Methods). Each resulting PASA “assembly”^36^ represents a distinct alternative isoform, and overlapping “assemblies” constitute a transcriptional unit (Supplementary Table 8). We identified 30,463 and 22,531 alternative isoforms comprising 24,947 and 19,750 transcriptional units in CCMP1383 and CCMP2088, respectively. When focusing only on DinoSL-type transcripts, these numbers are 8,714 and 6,576, comprising 7,146 and 5,110 transcriptional units respectively (Supplementary Table 8). In both isolates, alternative exons are the most common events observed among all transcript isoforms (e.g. 45.85% of all inferred events in CCMP2088), followed by alternative donor (22.05%) and acceptor (20.43%) sites (Supplementary Table 9).

The addition of DinoSL sequences was proposed to be a mechanism to split polycistronic pre-mRNA into monocistronic mature mRNA^29^. A little over one-half (50.45% in CCMP1383, 58.78% in CCMP2088) of these DinoSL-type transcriptional units are located within 5 kbp of one another (Fig. 2c). Interestingly, among the DinoSL-type transcript isoforms, the two most-enriched Pfam domains in both isolates are bacteriorhodopsin-like protein (PF01036) and cold-shock DNA-binding (PF00313) (Supplementary Table 10). To further assess the functional diversity of DinoSL-type transcripts, we sequenced 747,959 full-length transcripts from CCMP1383 specifically selected for DinoSL (Supplementary Table 6; see Methods). These transcripts comprised only 1,187 isoforms (3.9% of the total 30,463 isoforms; Supplementary Table 8). Similar functions are prevalent among these genes (Supplementary Table 10), thus lending support to our observation of functional bias in DinoSL-type transcripts. In addition, the frequency at which DinoSL-type transcripts are integrated back into the genome is likely dependent on their relative abundance in the nucleus. Therefore, transcripts containing relic DinoSLs are likely to be, or have been, highly expressed. In both isolates, the ice-binding (DUF3494) and the bacteriorhodopsin-like protein domains, both important for adaptation to cold (see below), are among the most-enriched features in transcripts encoded with a relic DinoSL.

### Prediction of gene models in *Polarella glacialis* is likely impacted by RNA editing

Using a gene-prediction workflow customised for dinoflagellate genomes^37^ (see Methods), we predicted 58,232 and 51,713 protein-coding genes in the CCMP1383 and CCMP2088 genomes, respectively (Table 2 and Supplementary Table 11). Of the 58,232 genes predicted in CCMP1383, 51,640 (88.68%) of the encoded proteins were recovered in CCMP2088 (Fig. 3a). Likewise, of the 51,713 genes predicted in CCMP2088, 46,228 (89.39%) of the encoded proteins were recovered in CCMP1383 (Fig. 3a). The difference in numbers of predicted genes and sequence dissimilarity observed between the two genomes could be explained in part by the presence of distinct transcript isoforms. Although transcriptome evidence can improve the quality of predicted genes, our results indicate that this evidence can also complicate prediction when a gene has multiple isoforms, more so when these isoforms are recovered unevenly between the two isolates. The generation of alternatively spliced mRNAs is likely explained by local adaptation in the polar regions that drives functional diversification.

**Fig. 3.**
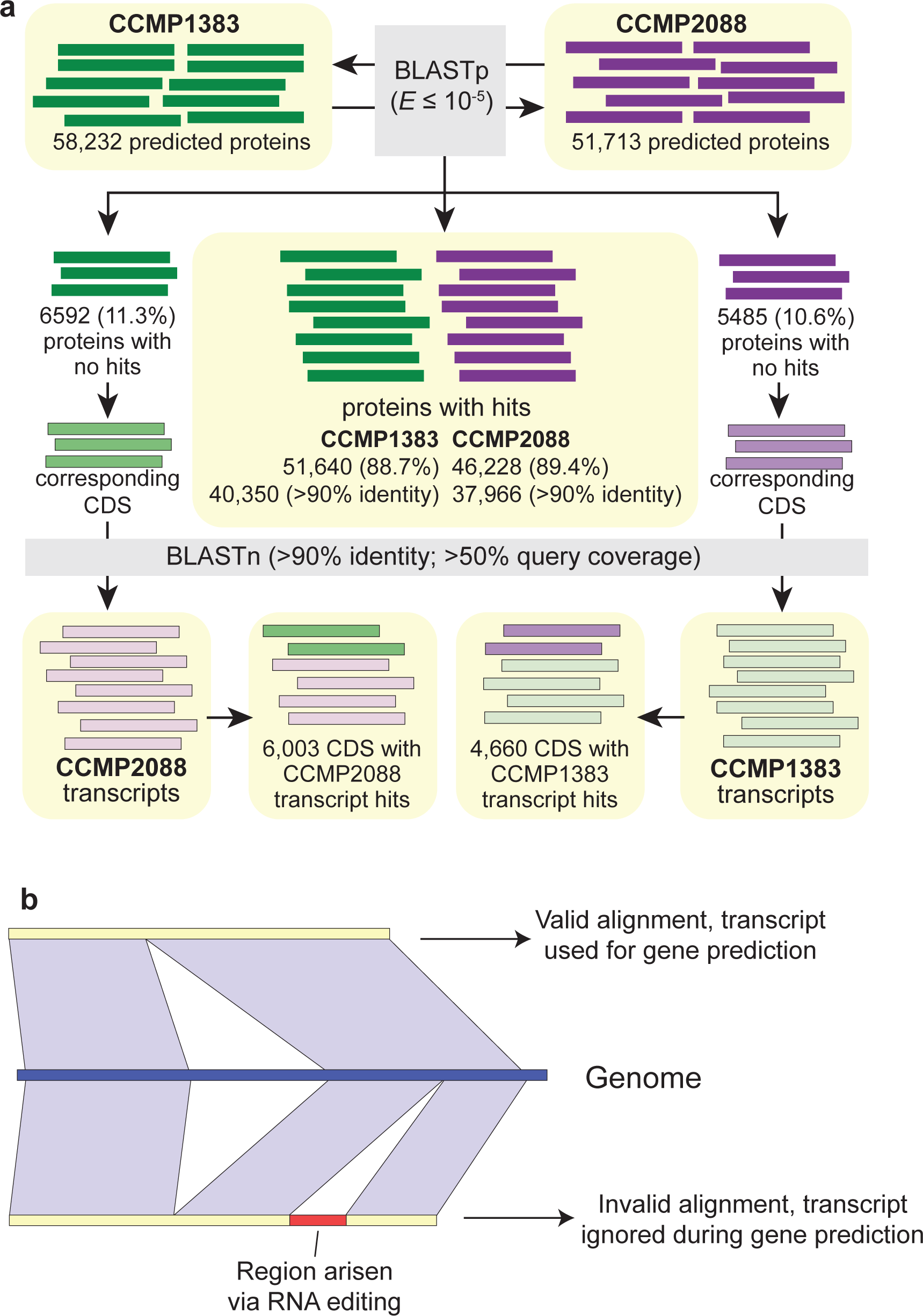
Comparison of predicted gene models between the two *P. glacialis* genomes. (a) The comparison of predicted proteins in CCMP1383 against those in CCMP2088 is shown, incorporating evidence from the corresponding transcriptome data. (b) Scenario of RNA editing (insertion/deletion) that would disrupt the alignment of a transcript to the genome.

**Table 2:** Predicted gene models in P. glacialis compared to representative publicly available dinoflagellate genomes. A more-comprehensive summary including gene models from all available dinoflagellate genomes is shown in Supplementary Table 11. Genes from the Symbiodiniaceae taxa are based on the revised predictions from Chen et al.^37^.

|  | *Polarella glacialis* |  | Symbiodiniaceae |  |  |  | Parasitic |
| --- | --- | --- | --- | --- | --- | --- | --- |
|  | CCMP1383 | CCMP2088 | *Symbiodinium microadriaticum* | *Breviolum minutum* | *Cladocopium goreau* | *Fugacium kawagutii* | *Amoebophrya ceratii* |
| **Genes** |  |  |  |  |  |  |  |
| Number of genes | 58,232 | 51,713 | 29,728 | 32,803 | 39,006 | 31,520 | 19,925 |
| Gene models supported by transcriptome (%) | 94.0 | 94.3 | 79.2 | 89.4 | 76.5 | 69.0 | 24.4 |
| G+C content of CDS (%) | 57.84 | 57.78 | 57.43 | 51.18 | 54.23 | 54.19 | 60.77 |
| **Exons** |  |  |  |  |  |  |  |
| Number of exons per gene | 11.64 | 10.84 | 19.21 | 19.03 | 12.46 | 11.63 | 3.39 |
| Average length (bp) | 105.67 | 108.71 | 115.44 | 101.38 | 130.47 | 158.13 | 577.84 |
| Total length (Mb) | 71.60 | 60.94 | 65.92 | 63.30 | 63.42 | 57.96 | 39.08 |
| **Introns** |  |  |  |  |  |  |  |
| Number of genes with introns (%) | 73.79 | 75.60 | 95.70 | 93.74 | 96.00 | 96.00 | 71.35 |
| Average length (bp) | 1,408 | 1,296 | 387.92 | 451.33 | 593.53 | 658.41 | 337.11 |
| Total length (Mb) | 837.95 | 636.20 | 210.00 | 267.00 | 265.35 | 220.58 | 16.08 |
| **Intergenic regions** |  |  |  |  |  |  |  |
| Average length (bp) | 21,625 | 20,922 | 15,108 | 5,983 | 9,538 | 18,050 | 1,525 |

Note: *Hematodinium* sp. is not shown as no predicted genes were reported.

Interestingly, almost all predicted proteins not recovered in the proteins of the counterpart isolate (i.e. 6003 of 6592 in CCMP1383 and 4660 of 5485 in CCMP2088) were recovered in the transcriptome of the counterpart isolate (Fig. 3a). These results indicate that some transcriptome evidence was not incorporated in ∼10% of the predicted genes in each genome. We hypothesise that this is likely due to RNA editing in *P. glacialis* (Fig. 3b). RNA editing has been characterised in the nuclear-encoded genes of *Symbiodinium microadriaticum*^38^, as well as organellar genes of other dinoflagellates^39, 40^. RNA editing may introduce changes in the transcripts (e.g., base substitutions or indels) affecting the identification of open-reading frames (e.g., disruption by in-frame stop codons or correction of premature stop codons) in the genome sequences, and thus impacting prediction of gene models.

Given the diploid genome assembly for each isolate, we assume that the number of predicted genes would approximate twice the number expected in a haploid genome (e.g. 50,000 genes in a diploid assembly versus 25,000 in a haploid assembly). The workflow used to predict genes in the Symbiodiniaceae genomes is the same as used in this study^37^, making the predicted gene features more comparable. As expected, the number of predicted genes in all six (haploid) genomes of Symbiodiniaceae is roughly similar to the haploid gene number for the two *P. glacialis* genomes (Table 2 and Supplementary Table 11). The proportion of genes predicted in *P. glacialis* that are supported by transcriptome evidence (∼94% for each isolate; Table 2) is much higher than in the Symbiodiniaceae isolates (∼79% averaged among six genomes^37^). This result may be explained by the more-extensive transcriptome data we generated in this study (using both RNA-Seq short-read and Iso-Seq full-length transcripts) to guide our gene-prediction workflow (see Methods), compared to the transcriptome data (based on RNA-Seq short-reads) available for the other isolates.

### Unidirectional tandem single-exonic genes in *P. glacialis*

In *P. glacialis*, the longer intergenic regions are largely comprised of repeats (Supplementary Fig. 3). Roughly a third of the intergenic regions (35.86% in CCMP1383; 34.97% in CCMP2088) are ≤ 5 Kbp in length. The fraction of these regions covered by repetitive elements is 32.92% (CCMP1838) and 32.93% (CCMP2088; Supplementary Fig. 3); these numbers are 59.65% and 59.05% (Supplementary Fig. 3) among intergenic regions >5 kbp. This observation suggests that expansion of repeats is greater in (and likely contributes to) longer intergenic regions in the genome. Approximately 50% of the analysed genes (26,580 in CCMP1383, 21,376 in CCMP2088) appear to have intergenic regions ≤ 5 kbp (Fig. 4a), indicating a tendency for these genes to occur in clusters. Adjacent genes were clustered if their intergenic regions were < 5 Kbp and clusters were considered unidirectional if all genes in that cluster were encoded in the same direction. Remarkably, almost all of these clustered genes (24,276 and 19,544 respectively for those of CCMP1383 and CCMP2088; ∼40% of total genes in each genome) are encoded unidirectionally. These unidirectional gene clusters may represent a mechanism in *P. glacialis* (and potentially the order Suessiales) to ensure transcriptional efficiency, with genes in close physical proximity potentially transcribed together. In some cases, these gene clusters encode the same or similar functions; e.g., 22 unidirectionally encoded genes in CCMP1383 (and 19 in CCMP2088) putatively encode the major basic nuclear protein 2.

**Fig. 4.**
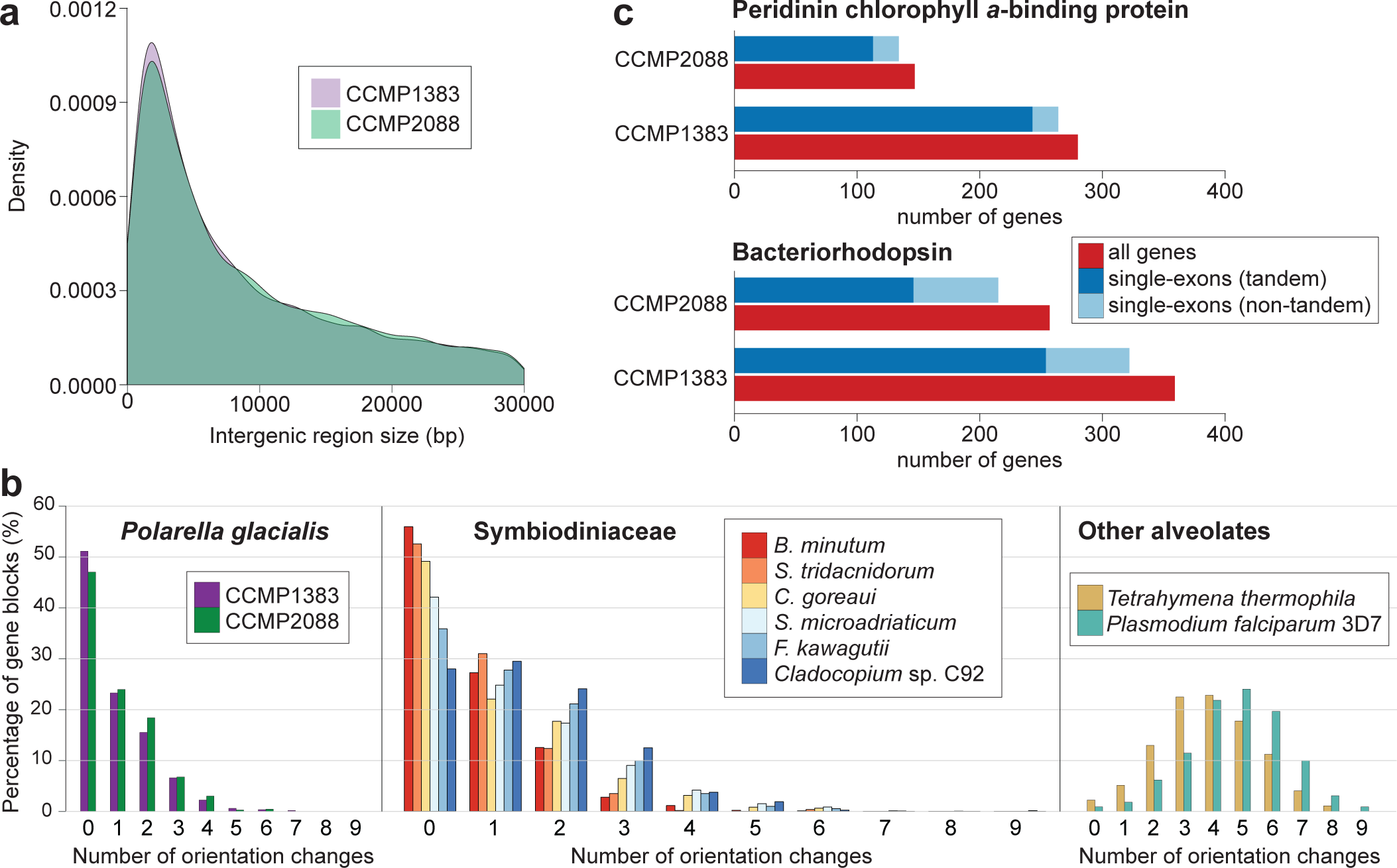
Intergenic regions and tandemly repeated genes. (a) Distribution of the sizes of intergenic regions (in bp; ≤ 30,000 bp) shown for the assembled *P. glacialis* genomes of CCMP1383 and CCMP2088. (b) Frequency of strand orientation changes in ten-gene windows generated from the predicted genes from isolates of *P. glacialis*, Symbiodiniaceae, and the other alveolates of *Tetrahymena thermophilia* (ciliate) and *Plasmodium falciparum* 3D7 (apicomplexan). (c) Number of tandemly repeated and/or single-exonic genes in CCMP1383 and CCMP2088, shown for genes encoding bacteriorhodopsin and peridinin chlorophyll *a*-binding proteins.

The small number of strand-orientation changes within ten-gene windows was used in the genome study of *B. minutum*^14^ to illustrate the tendency for genes to be encoded unidirectionally. Using the same approach, *P. glacialis* and other Symbiodiniaceae taxa were found to encode genes unidirectionally, with most ten-gene windows having no more than three strand-orientation changes (Fig. 4b). In contrast, this pattern was not observed in the genomes of other alveolates (i.e., the ciliate *Tetrahymena thermophilia* and the apicomplexan *Plasmodium falciparum* 3D7), for which most ten-gene windows have three to six strand-orientation changes (Fig. 4b). Whereas this analysis does not consider the distance between genes, the unidirectionality of coding regions reflects a feature common within the order Suessiales (and potentially all dinoflagellates).

Among the predicted genes in both genomes, 4,898 (CCMP1383; 8.4%) and 3,359 (CCMP2088; 6.5%) are located (nested) within introns of multi-exon genes. Although most cases (71.02% in CCMP1383 and 74.90% in CCMP2088) represent one nested gene per multi-exon gene, in extreme cases, we observed 18 (CCMP1383) and 24 (CCMP2088). Supplementary Fig. 4 shows an example of 15 nested genes of CCMP1383 spanning three introns of the gene putatively encoding an alanine–tRNA ligase. Among the nested genes within each intron, five encode fucoxanthin chlorophyll *a/c*-binding protein, and four, light-harvesting complex protein. The validity of the nested gene structure was confirmed by expression evidence based on full-length transcripts.

Of particular interest, we recovered 15,263 (26.2%) and 12,619 (24.4%) single-exon genes in CCMP1383 and CCMP2088 respectively (Table 2). These proportions are higher than those in symbiodiniacean genomes (< 12% of genes; Table 2 and Supplementary Table 11). Of all recovered single-exon genes, >80% are recovered in both isolates, and almost all (99.08% of those in CCMP1383, 98.12% of those in CCMP2088) are supported by transcriptome evidence (including full-length transcripts that were selected for 5′-cap and 3′-polyadenylation sites). These results suggest that these genes are *bona fide P. glacialis* genes (i.e., not bacterial contaminants, nor artefacts of our gene-prediction workflow). Many of the Pfam domains enriched in the single-exon genes are also enriched in the predicted genes of *P. glacialis* compared with symbiodiniacean genes (see also below; Supplementary Table 12). Enriched features of *P. glacialis* such as bacteriorhodopsin-like protein (PF01036), peridinin-chlorophyll *a*-binding (PF02429), and DUF3494 (PF11999) are encoded as single-exon genes. A number of other domains are enriched in the single-exon genes in both *P. glacialis* isolates. The bacterial DNA-binding protein domain (PF00216), which is predominantly found in bacteria, is enriched and potentially has arisen in *P. glacialis via* lateral genetic transfer. The reverse transcriptase (PF00078) domain is also enriched and is likely involved in the activity of retrotransposons in the *P. glacialis* genomes.

### What makes Polarella Polarella?

We compared the annotated functions of *P. glacialis* genes against those from other Symbiodiniaceae genomes^12–15^. When comparing the annotated Pfam domains, we observed a significant over-representation of DUF3494 (PF11999), cold-shock (PF00313), and chlorophyll *a-b* binding (PF00504) domains in *P. glacialis* relative to Symbiodiniaceae (Supplementary Table 13), as we previously observed in an independent transcriptome analysis^41^. In this study using high-quality gene models predicted from genome data, we also observed over-representation (adjusted *p*-value ≤ 10^-40^) of pentatricopeptide repeat (PF13041, PF13812, PF01535), ATP synthase subunit C (PF00137), bacteriorhodopsin-like protein (PF01036), and peridinin-chlorophyll *a*-binding (PF02429) in *P. glacialis*. Interestingly, the peridinin-chlorophyll *a*-binding and bacteriorhodopsin-like domains are predominantly encoded in blocks of tandemly repeated single-exon genes, implicating hundreds of genes in *P. glacialis* (Fig. 4c). All but one of these gene blocks (i.e., a contiguous region containing two or more genes) are unidirectionally encoded. Within each isolate, some of these features (e.g., peridinin-chlorophyll *a*-binding and bacteriorhodopsin-like domains) were also enriched among the tandemly repeated genes in *P. glacialis* (when compared against all genes within each *P. glacialis* isolate; Supplementary Table 14). Peridinin-chlorophyll *a*-binding protein was thought to be encoded as 5000 single-exon gene copies in tandem repeat blocks in the bloom-forming dinoflagellate *Lingulodinium polyedra*^21^, and these coding genes are thought to be monocistronic^42^. This protein may be universally important in free-living dinoflagellates, and potentially to a lesser extent among symbiotic lineages of these algae^43^.

The tendency of tandemly repeated genes to have fewer introns was also reported in the bloom-forming *Amphidinium carterae*^44^. In combination with our other results (above), our observations suggest gene-family expansion through tandem duplication drives the genome evolution of *P. glacialis*, and potentially of other free-living dinoflagellates. The use of a DinoSL sequence to split polycistronic transcripts into mature RNAs^29^ may facilitate this mechanism.

Bacterial-derived rhodopsin, a transmembrane protein involved in bacterial phototrophy independent of chlorophyll through retinal binding, is encoded in diverse dinoflagellate lineages^45, 46^. The proton-pump type rhodopsins can create a proton gradient to drive synthesis of ATPase, in lieu of photosynthesis^47, 48^. An earlier gene-expression analysis of the bloom-forming *Prorocentrum donghaiense*^49^ revealed that proton-pump rhodopsins may compensate for photosynthesis under light-deprived conditions. These rhodopsins were also found to be highly expressed in diatoms under iron-deficient conditions^50^. All genes in both *P. glacialis* isolates have top hits to the sequences encoding proton-pump rhodopsins in *Oxyrrhis marina*. These rhodopsins were previously found to be more abundantly expressed in *O. marina* than the sensory-type rhodopsins involved in light-harvesting for photosynthesis^51^. We hypothesise that the over-representation of rhodopsin and other photosynthesis-related genes in *P. glacialis* is an adaptation to light-limited (and potentially iron-limited) conditions, as expected in the ice-brine channels were one of the samples was collected^10^.

A previous study based on transcriptome analysis^41^ revealed “dark” proteins (i.e., they lack an annotation based on sequence-similarity searches against characterised proteins; see Methods) that are conserved and/or lineage-specific in dinoflagellates. Using 302,231 protein sequences predicted from genome data of *P. glacialis* and six Symbiodiniaceae species (Supplementary Table 15), we constructed 35,751 putatively homologous protein sets that consist of 85.3% of the total protein sequences analysed. Of these sets, 8,673 (24.26%) containing 10.63% of the clustered proteins (27,425 proteins; 9.07% of 302,231) were classified as dark. The number of dark proteins (and hence dark genes) from each dataset (Supplementary Table 15) was largely congruent with the proportions of dark genes reported previously^41^. Of the 8,673 dark homologous sets, 4,602 (53.06%) contain sequences from only *P. glacialis*; 4,540 (98.65% of the 4602) contain sequences from both isolates, so they are unlikely to have arisen due to assembly artefacts. We consider a dark set as single-exonic if all its members are encoded in single exons, and a dark set as multi-exonic if at least one member is encoded in multiple exons. Following this definition, most (3,149; 68.43%) of the 4602 *P. glacialis*-specific sets are multi-exonic, whereas 1,453 (31.57%) are single-exonic. Of the 1,453 single-exonic dark sets, 714 (49.14%) are supported by IsoSeq data and 1,449 (99.72%) by IsoSeq and/or RNA-Seq data. Therefore, these genes likely represent true genetic (and functional) innovation specific to *P. glacialis*.

### Single evolutionary origin of ice-binding domains in dinoflagellates

The Pfam domain DUF3494, a known ice-binding domain^52^, is over-represented in cold-adapted dinoflagellates^41^. In both *P. glacialis* isolates (Supplementary Table 16), most putative ice-binding genes encode only the DUF3494 domain. They are encoded in single exons, and in unidirectional, tandemly repeated blocks, potentially as a mechanism to enhance the efficiency of gene expression. Because the DUF3494 domain in many species has arisen *via* lateral genetic transfer^52^, the presence of these genes in this configuration suggests that they might have arisen *via* the same mechanism in *P. glacialis*.

Fig. 5 shows part of a phylogenetic tree reconstructed with 1,080 sequences of available DUF3494 domains encompassing Archaea, Bacteria, and eukaryotes; the complete tree is available as Supplementary Data 1. All DUF3493 sequences from the dinoflagellates (*P. glacialis*, *Heterocapsa arctica*, *Scrippsiella hangoei* and *Peridinium aciculiferum*), plus some sequences from the ice diatom *Fragilariopsis cylindrus*, form a strongly supported clade (bootstrap support [BS] = 100% based on ultrafast bootstrap approximation^53^) (Fig. 5).

**Fig. 5.**
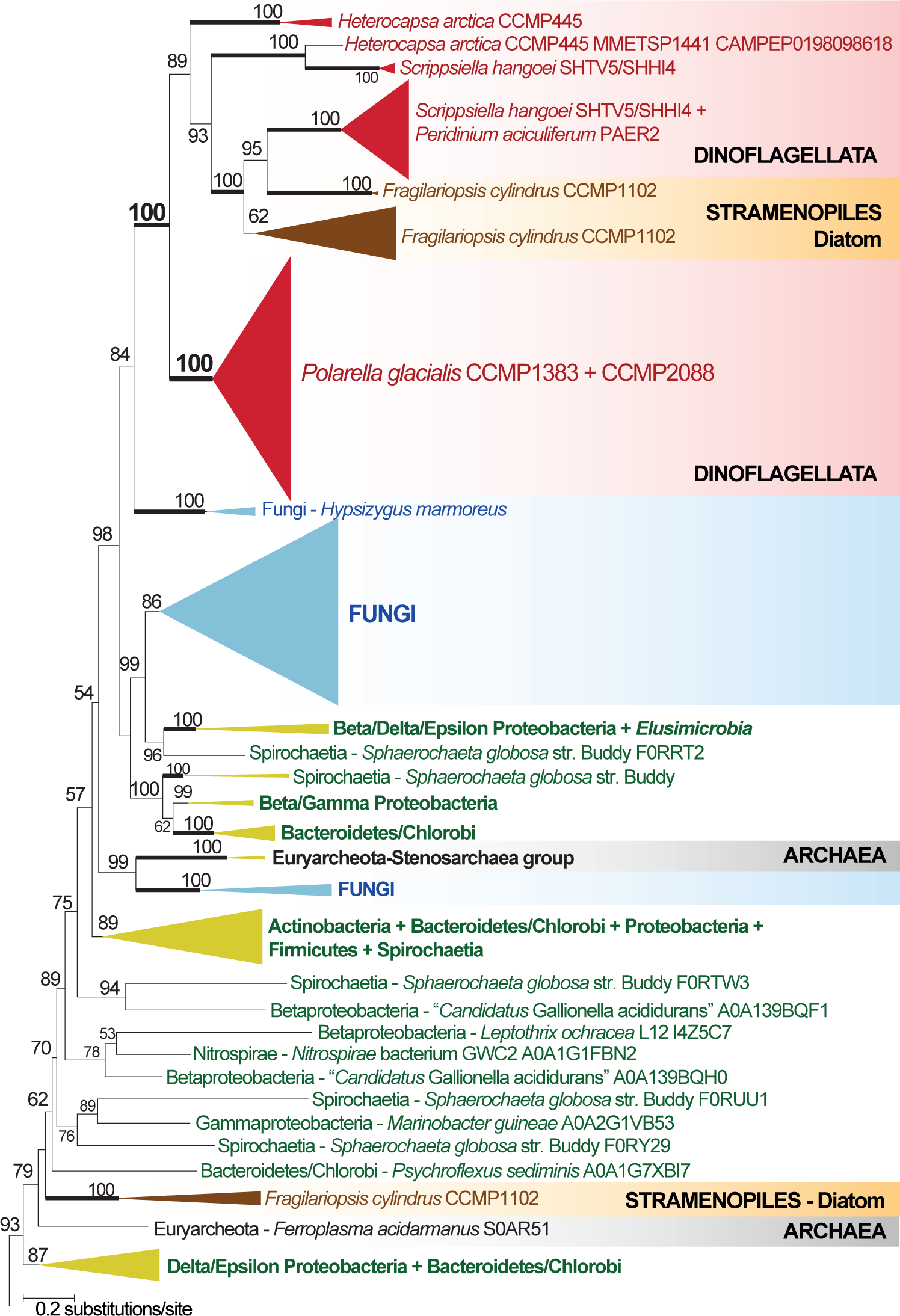
Evolutionary history of ice-binding domains in *P. glacialis* and dinoflagellates. Only a small part of the 1,080-taxon maximum likelihood protein tree is shown. Support values, based on 2,000 ultrafast bootstrap approximations, are shown at the internal nodes. Only values >50% are shown. The unit of branch length is the number of substitutions per site.

Within this dinoflagellate + diatom clade, the 169 DUF3494 sequences from *P. glacialis* (97 from CCMP1383, 72 from CCMP2088) form a strongly supported monophyletic clade (BS 100%), indicating that these domains in *P. glacialis* have an evolutionary history that is distinct from other dinoflagellates. In comparison, the domains in the ice diatom *F. cylindrus* were recovered in three distinct clades on the tree (two shown in Fig. 5), indicating their independent origins. As previously reported, DUF3494 domains in eukaryotes trace their origins to multiple events of lateral genetic transfer from bacteria and other eukaryotes^54, 55^. We also observed this pattern in our phylogenetic analysis although the origin of these domains in dinoflagellates remains unclear, with potential sources being Proteobacteria, Bacteroidetes/Chlorobi, or Euryarchaeota that also gave rise to the domains in some fungal species. Fungi are also distributed in multiple clades on this tree (Supplementary Data 1). The DUF3494 domains we recovered from the bacterial genomes (see Methods and Supplementary Note) were grouped with their closely related species within the corresponding phylum (i.e. Proteobacteria and Bacteroidetes/Chlorobi) in distinct clades, indicating that they are indeed prokaryotic. These results indicate that all ice-binding domains in dinoflagellates share a single common origin likely from a Proteobacteria or Bacteroidetes/Chlorobi source, and that those specific to *P. glacialis* have a distinct evolutionary history that may reflect niche specialisation.

## Discussion

We generated two draft *de novo* diploid assemblies of *P. glacialis*, the first of any free-living psychrophilic dinoflagellates, and high-quality gene models supported by full-length transcriptomes. Genome features of *P. glacialis* (Fig. 6) elucidate how the genomes of dinoflagellates have evolved to adapt in a harsh environment. The difference in genome sizes between the two isolates highlights the extensive structural divergence of genomes within a dinoflagellate species. The abundance of repetitive elements and LTRs in the genomes suggests their important role in shaping the genome evolution of these isolates, potentially contributing to the genome-size difference. The molecular mechanisms and selective pressure that contribute to the larger genome size in the Antarctic *versus* the Arctic isolate remains an open question, and can best be addressed using assembled genomes at chromosomal resolution. The trans-spliced DinoSL was thought to be a global signature of all transcripts in dinoflagellates, but our results reveal only a small proportion of full-length transcripts encode DinoSL (and with relic DinoSLs) and remarkably, these transcripts mostly encode functions that are critical for adaptation to cold and to low-light conditions, both relevant to the natural habitat of *P. glacialis* in ice-brine channels. In addition, genes encoding these functions are unidirectionally encoded, often in a tandemly repeated single-exonic structure. This distinctive gene organisation is likely a feature of free-living dinoflagellates, and may serve to enhance transcriptional efficiency of critical functions. The independently evolved ice-binding domains and the lineage-specific dark genes in *P. glacialis* highlight functional innovation in dinoflagellate genomes relevant to environmental adaptation and niche specialisation as successful psychrophiles in the extreme cold environment.

**Fig. 6.**
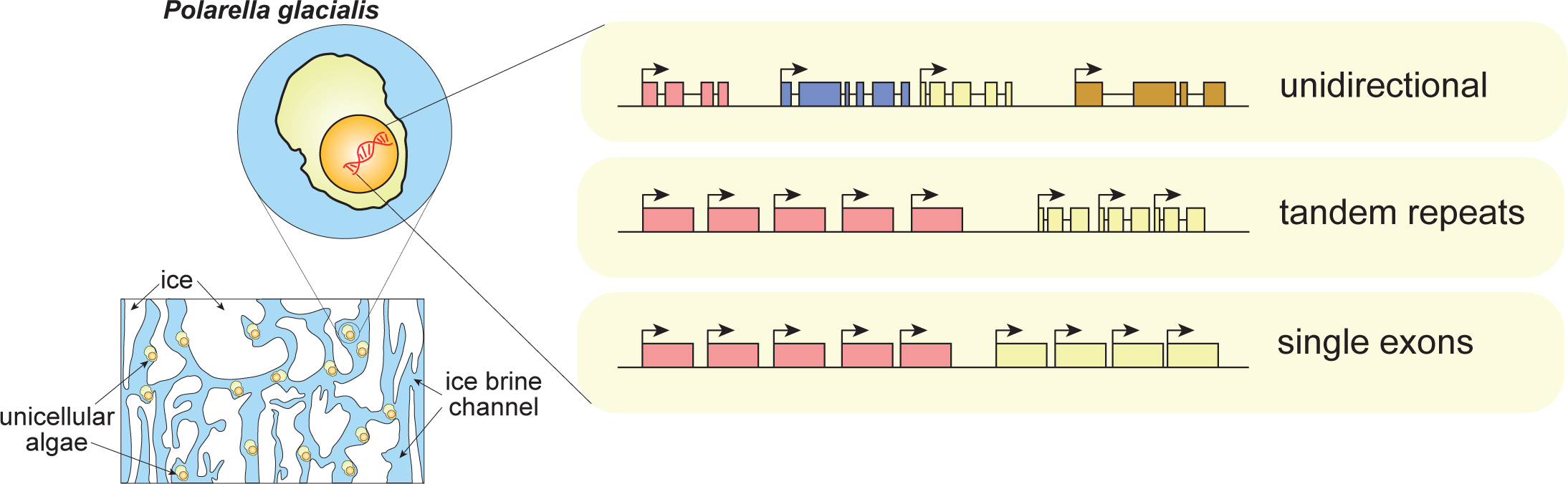
Genome features of *Polarella glacialis* as a psychrophilic, free-living dinoflagellate. Summary of key genome features of *P. glacialis*, focusing on unidirectionality of coding genes, tandemly repeated genes, and single-exon genes.

## Methods

### Cultures of Polarella glacialis

The cultures of *Polarella glacialis* isolates were acquired from the National Center for Marine Algae and Microbiota at the Bigelow Laboratory for Ocean Sciences, Maine, USA. Both cultures were maintained in f/2 medium without silica^56^ (100mL culture in 250mL conical flasks, 12h:12h light:dark cycle, 90 µmol photon·m^-2^·s^-1^, 4°C). The cultures were treated with ampicillin (100 μg·mL^−1^), kanamycin (50 μg·mL^−1^) and streptomycin (50 μg·mL^−1^) for 24 hours before cell harvest. For extraction of nucleic acids, the cells (50mL; >10^6^ per mL) were harvested by centrifugation (3000 *g*, 5 min). The resulting cell pellet was rinsed with 0.22μm-filtered artificial seawater (Instant Ocean salt mixture, 33.3 g.L^-1^; 1 mL), transferred to an 1.5mL-tube, and collected by further centrifugation (3000 *g*, 5 min). The supernatant (seawater) was removed, and the tube was immediately snap-frozen with liquid nitrogen and stored at -80°C until DNA/RNA extraction.

### Extraction of genomic DNA and total RNA

Genomic DNA was extracted following the 2xCTAB protocol with modifications. The cells were suspended in a lysis extraction buffer (400 μL; 100 mM Tris-Cl pH 8, 20 mM EDTA pH 8, 1.4 M NaCl), before silica beads were added. In a freeze-thaw cycle, the mixture was vortexed at high speed (2 min), and immediately snap-frozen in liquid nitrogen; the cycle was repeated 5 times. The final volume of the mixture was made up to 2% w/v CTAB (from 10% w/v CTAB stock; kept at 37 °C). The mixture was treated with RNAse A (Invitrogen; final concentration 20 μg/mL) at 37°C (30 min), and Proteinase K (final concentration 120 μg/mL) at 65°C (2 h). The lysate was then subjected to standard extractions using equal volumes of phenol:chloroform:isoamyl alcohol (25:24:1 v/v; centrifugation at 14,000 g, 5 min, RT), and chloroform:isoamyl alcohol (24:1 v/w; centrifugation at 14,000 g, 5 min, RT). DNA was precipitated using pre-chilled isopropanol (gentle inversions of the tube, centrifugation at 18,000 *g*, 15 min, 4 °C). The resulting pellet was washed with pre-chilled ethanol (70% v/v), before stored in Tris-HCl (100 mM, pH 8) buffer. Total RNA was extracted using RNeasy Plant Mini Kit (Qiagen) following the manufacturer’s protocol. The concentration of DNA or RNA was determined with NanoDrop (Thermo Scientific), and a sample with A230:260:280 ≈ 1.0:2.0:1.0 was considered appropriate for sequencing.

### Generation of genome data

For generation of short-read sequence data, samples of genomic DNA were sequenced using Illumina HiSeq2500 (Australian Genome Research Facility, Melbourne), and HiSeq4000 (Translational Research Institute and the Australian Genome Research Facility, Brisbane) platforms (Supplementary Table 17). For each isolate, two paired-end TruSeq libraries (inserts of ∼250bp and 600bp for HiSeq2500; ∼350bp and ∼600bp for HiSeq4000), and three mate-pair Nextera libraries (inserts of ∼2, 5 and 10Kb) were generated for sequencing (in 2x150 bases). In total, we generated 399.3 Gbp of Illumina short-read sequencing data for CCMP1383, and 746.7 Gbp for CCMP2088 (Supplementary Table 17).

For generation of long-read data, samples of genomic DNA were sequenced using the PacBio Sequel platform available at the Queensland University of Technology Central Analytical Research Facility, and at the Ramaciotti Centre of Genomics (University of New South Wales, Sydney). In total, 15 SMRT cells were sequenced for CCMP1383 producing 9.3 million subreads (74.6 Gbp), and 7 SMRT cells for CCMP2088 generating 4 million subreads (35.9 Gbp); see Supplementary Table 2 for details.

### Generation of transcriptome data (RNA-Seq)

For generation of RNA-Seq data, total RNA samples were sequenced at the Australian Genome Research Facility (Brisbane) using the Illumina HiSeq4000 platform. Illumina paired-end (2x150 bp reads) RNA-Seq data was generated for both CCMP1383 (55.4 Gbp) and CCMP2088 (61.7 Gbp); see Supplementary Table 18 for details.

### Generation of full-length transcript data (PacBio IsoSeq)

Using the extracted total RNA samples (above), a full-length cDNA library was constructed for each of CCMP1383 and CCMP2088 using the TeloPrime Full-Length cDNA Amplification Kit (Lexogen, Vienna) following the kit manual. Two cDNA synthesis reactions were carried out in parallel for each sample, with 2 μg of total RNA used as starting material in each reaction. Double-stranded cDNA resulting from the two reactions was combined before performing PCR amplification using the TeloPrime PCR Add-on Kit (Lexogen, Vienna). For each sample, 16 parallel PCRs were carried out using 22 amplification cycles and 2 μL of double-stranded cDNA per reaction as template; the PCR products were pooled together and then split into two fractions, which were purified using 1x and 0.5x AMPure PB beads (Pacific Biosciences, California), respectively, and pooled at equal molarity. For sample CCMP1383, a total of 2.52 μg of purified full-length cDNA was obtained and was used for PacBio SMRTbell library preparation with the SMRTbell Template Prep Kit 1.0 (Pacific Biosciences, California); the library was sequenced on 4 SMRT cells v2 LR using 20-hour movies on a Sequel platform at the Institute for Molecular Bioscience Sequencing Facility (University of Queensland, Brisbane). For CCMP2088, 1.95 ng of cDNA were obtained and submitted to The Ramaciotti Centre for Genomics (University of New South Wales, Sydney) for SMRTbell library preparation and sequencing on a PacBio Sequel System, also using 4 SMRT Cells v2 LR and 20-hour movies.

In addition to the cDNA libraries described above, a spliced-leader-specific transcript library was generated for CCMP1383. Four parallel PCR reactions were performed with the TeloPrime PCR Add-on Kit (Lexogen, Vienna) using 12 amplification cycles, conserved spliced leader fragment (5′-CCGTAGCCATTTTGGCTCAAG-3′) as forward primer, TeloPrime PCR 3′ primer as reverse primer, and 2 μL of double-stranded cDNA synthesised using the TeloPrime Full-Length cDNA Amplification Kit (Lexogen, Vienna) as template.

The PCR products were pooled and purified (same method as above), resulting in 987 ng of cDNA. SMRTbell library construction was carried out using the PacBio SMRTbell Template Prep Kit 1.0, followed by sequencing on 2 SMRT cells v2 LR using 20-hour movies on the Sequel at the Institute for Molecular Bioscience Sequencing Facility (University of Queensland, Brisbane). Data yield from each SMRT cell is detailed in Supplementary Table 19. All genome and transcriptome sequencing data are available at NCBI GenBank *via* BioProject accession PRJEB33539.

### Processing of sequence data

Adaptor sequences were removed and low quality bases trimmed from paired-end reads using Trimmomatic v0.35 (LEADING:10 TRAILING:10 SLIDINGWINDOW:4:30 MINLEN:50)^57^, overlapping read pairs (250 bp insert size) were merged using FLASH v1.2.11 (max-overlap 85)^58^. Mate-pair reads were processed using the NextClip v1.3 pipeline^59^ using the preliminary CLC assembly as a reference. Only the category A, B and C mate-pair reads were retained from the NextClip analysis. Further trimming of the mate-pair reads to remove low quality regions and adapters was performed using Trimmomatic v0.35 (LEADING:10 TRAILING:10 SLIDINGWINDOW:4:20 MINLEN:25).

Trimmed paired-end reads were mapped using bowtie2^60^ against the initial CLC assembly and the mean insert size and standard deviation were computed using Picard tools v2.6.0 “CollectInsertSizeMetrics”. Mate-pair sequences were aligned only against scaffolds from the initial assembly with a length >15 Kbp using bbmap (rcs=f pairedonly=t ambig=toss). The different approach taken for the mate-pair reads was to discard ambiguously mapped reads (i.e. reads that map equally well to multiple locations), as they have a more pronounced effect on inset size estimation with mate-pair data. The maximum insert size set during the alignment stage of both mate-pair and paired-end libraries was double the maximum expected insert size.

Illumina RNA-Seq data was trimmed for adapters and low-quality regions using Trimmomatic v0.35 (LEADING:10 TRAILING:10 SLIDINGWINDOW:4:30 MINLEN:50).

PacBio IsoSeq data from each SMRT cell was polished (--polish) using the circular consensus sequencing tool (ccs v3.1.0 https://github.com/PacificBiosciences/unanimity/blob/develop/doc/PBCCS.md); only polished reads with a quality >0.99 were retained. Primers were removed from the polished reads using lima v1.8.0 (https://github.com/pacificbiosciences/barcoding) in IsoSeq mode.

Only reads with the correct 5-prime/3-prime primer configuration were retained. The PacBio IsoSeq tool v3.1.0 was used to remove concatemers (using the refine option) from the primer trimmed reads (https://github.com/PacificBiosciences/IsoSeq3/blob/master/README_v3.1.md).

### Genome-size estimation using *k*-mers

The *k*-mers frequency distribution in the trimmed paired-end reads (including merged) was used to estimate genome size and to assess ploidy. Genome size estimation was conducted following the approach described in Liu et al.^12^. The enumeration of *k*-mers was performed using Jellyfish^61^ at *k* = 17, 19, 21, 23, 25, 27, 29 and 31. For diploid genomes, a bimodal *k*-mer-count distribution is expected, genome size estimated from the first peak represents the diploid state, and that estimated from the second peak represents the haploid state. The standard theoretical model of a diploid genome in GenomeScope^62^ was used (*k*-mer size 21) to verify the diploidy observed in the sequence data (Supplementary Fig. 1).

### *De novo* genome assemblies

Initial *de novo* assemblies for CCMP1383 and CCMP2088 were generated independently using CLC Genomics Workbench (v7.5) (default parameters), incorporating all trimmed paired-end reads (using merged reads where applicable). The initial CLC assemblies was further processed using the Redundans package (retrieved 10 March 2019)^63^ using the trimmed paired-end and mate-pair reads.

Final genome assemblies for each isolate were generated with MaSuRCA v3.2.8^64^ using untrimmed paired-end reads, trimmed mate-pair reads, and PacBio reads (>5 Kbp). For each isolate, the parameters of estimated assembly size in MaSuRCA was set based on the average estimated haploid genome size (in Supplementary Table 3), ploidy was set to two. Scaffolds <1 Kbp were discarded from the final assembly. The MaSuRCA assembler will attempt to remove redundant and (when ploidy set to two) homologous scaffolds, producing a haploid assembly. For both isolates, this step was ineffective in identifying and collapsing homologous scaffolds into a haploid assembly.

Trimmed and merged paired-end reads were mapped using bowtie2^60^ against the MaSuRCA diploid assembly and the purge_haplotigs^65^ program (v1.0.2; ‘purge_haplotigs contigcov -l 5-m 63 -h 120’ for CCMP1383 and ‘purge_haplotigs contigcov -l 40 -m 102 -h 170’ for CCMP2088) was run in an attempt to reconstruct a haploid assembly for each isolate. Unfortunately, purge_haplotigs was unable to fully reduce the CCMP1383 and CCMP2088 assemblies into haploid representations. As no reliable haploid assembly could be generated, the diploid assemblies generated by MaSuRCA were used for downstream analysis.

### Identification and removal of archaeal, bacterial, and viral sequences

Identification and removal of contaminant sequences in the genome assemblies was assessed using a method similar to that of Aranda et al.^13^. Genome scaffolds were compared using BLASTN against a database of archaeal, bacterial and viral genome sequences retrieved from RefSeq (release 88). Scaffolds were retained, and considered non-contaminant, if ≤10% of their length was covered by BLAST hits with a bit score >1000 and *E* ≤ 10^-20^. Further identification and removal of contaminant sequences was conducted following Chen et al.^37^. Scaffolds were removed if they had a G+C content and length outside the expected normal distribution; no putative contaminant scaffolds were identified using this approach. Statistics of the putative contaminant sequences are presented in Supplementary Table 20. Genes were predicted in the putative contaminant sequences using PROKKA^66^ (v1.13.3; --metagenome) and annotated with protein domains identified using pfam_scan.pl (v1.6; Pfam database release 31) at *E*-value < 0.001 following earlier studies^15, 41, 67^.

### Identification and removal of organelle sequences

The coding sequences from the plastid genome of *Cladocopium* sp. C3 (formerly *Symbiodinium* subtype C3) were used to identify putative plastid sequences from the assembly^68^. A scaffold was considered to be putatively of plastid origin if it shared significant sequence similarity (BLASTN) to one of the above sequences, covering >75% the sequence length at *E* ≤ 10^−10^.

The complete CDS of *cox1*, *cox3* and *cob* from *Breviolum minutum* (LC002801.1 and LC002802.1) were retrieved (because no complete sequences yet exist for *Polarella glacialis*) and used to identify putative mitochondrial scaffolds (Supplementary Table 21 and Supplementary Note). A scaffold was considered putative mitochondrial if it shared significant similarity (BLASTN, max_target_seqs 10000) to one of the above sequences, covering >75% of the sequence length at *E* ≤ 10^−10^. Statistics of the putative mitochondrial and plastidic sequences are presented in Supplementary Table 20.

### Customised gene prediction workflow tailored for dinoflagellate genomes

An *ab initio* gene prediction approach adapted from Liu et al.^12^ was applied to the genomes of *P. glacialis*; this approach was also reported in Chen et al.^37^. For each genome assembly, a *de novo* repeat library was first derived using RepeatModeler v1.0.11 (http://www.repeatmasker.org/RepeatModeler/). All repeats (including known repeats in RepeatMasker database release 20170127) were masked using RepeatMasker v4.0.7 (http://www.repeatmasker.org/).

We used transcriptome data generated in this study to guide gene prediction of assembled genomes. For RNA-Seq data, we assembled the reads using Trinity^69^ independently in “de novo” mode (v2.6.6) and “genome-guided” mode (v2.8.4). The combined Trinity assemblies were trimmed using SeqClean (https://sourceforge.net/projects/seqclean/). The RNA-Seq and polished PacBio IsoSeq transcripts were combined into gene assemblies using PASA v2.3.3^36^ that was customised (available at http://smic.reefgenomics.org/download/) to recognise an additional donor splice site (GA). TransDecoder v5.2.0^36^ was used to predict open reading frames on the PASA assembled transcripts. Complete proteins (CDS with both start and stop codons) predicted by TransDecoder that had valid genome coordinates and more than one exon were retained for further analysis.

These proteins were searched (BLASTP, *E* ≤ 10^−20^) against a customised protein database that consists of RefSeq proteins release 88 and other predicted Symbiodiniaceae and *Polarella* proteins (Supplementary Table 22). Only nearly full-length proteins were included in the subsequent analysis; we defined nearly full-length proteins as sequences with a BLAST hit that covered >80% of both the query and subject sequences.

The nearly full-length gene models were checked for TEs using HHblits v2.0.16^70^ (-p 80 -e 1E-5 -E 1E-5) searching against the JAMg transposon database (https://sourceforge.net/projects/jamg/files/databases/), as well as with Transposon-PSI (http://transposonpsi.sourceforge.net/). Gene models containing TEs were removed from the gene set, and redundancy reduction was conducted using CD-HIT v4.6.8^71^ (ID = 75%; -c 0.75-n 5). The remaining gene models were processed using the Prepare_golden_genes_for_predictors.pl (http://jamg.sourceforge.net/) script from the JAMg pipeline (altered to recognise GA donor splice sites). This script produces a set of “golden genes”, which were used as a training set for the gene-prediction tools AUGUSTUS v3.3.1^72^ and SNAP version 2006-07-28^73^. We used a customised code of AUGUSTUS (available at http://smic.reefgenomics.org/download/) so it recognises GA donor splice sites, and trained it to predict both coding sequences and untranslated regions; SNAP was trained for both GT and GC donor splice sites. Soft-masked genomes were passed to GeneMark-ES^74^ for training and gene prediction.

UniProt-SwissProt (retrieved 27/06/2018) proteins and other predicted Symbiodiniaceae and *Polarella* proteins (Supplementary Table 22) were combined to produce a set of gene models using MAKER v2.31.8 (altered to recognise GA donor splice sites)^75^ in protein2genome mode; the custom repeat library was used by RepeatMasker as part of MAKER prediction.

Two sets of predicted protein coding genes, one derived using the RNA-Seq data and one using the IsoSeq data, were constructed using PASA (--ALT_SPLICE -N 2) and TransDecoder (ORF prediction guided by Pfam database release 31). Gene models constructed using the IsoSeq data were assumed to be full-length and an extra step was taken to correct predicted proteins produced by TransDecoder that were five-prime partial. If a protein had an in-frame start codon within either the first 30 positions or the first 30% of the sequence, that position was then considered as the start of that sequence. Sequences not satisfying these criteria were left unchanged. A primary set of predicted genes was produced using EvidenceModeler v1.1.1^76^, which had been altered to recognise GA donor splice sites. This tool combined the gene models from PASA RNA-Seq, PASA IsoSeq (with corrected start positions where applicable), AUGUSTUS, MAKER protein2genome and GeneMark-ES, into a single set of evidence-based predictions. EvidenceModeler was allowed to predict genes within introns of other genes if the intron was >10,000 bp (--search_long_introns 10000).

Unlike Liu et al.^12^, we did not incorporate gene predictions from the SNAP program into the EvidenceModeler stage of the prediction workflow. This was done because SNAP produced an excessive number of overlapping genes that were on encoded on opposite strands. As genes encoded in this manner were not found to be supported in the transcriptome, we decided to exclude the results of this program from our predictions. We did not provide the location of putative repetitive elements to EvidenceModeler either, as multi-copy genes are often classified as repeats by RepeatModeler and would have been excluded from our final gene set. The weightings used for integration of gene models with EvidenceModeler were: PASA IsoSeq (with corrected start sites) 15, PASA RNA-Seq 10, Maker protein2genome 4, AUGUSTUS 1 and GeneMark-ES 1. EvidenceModeler gene models were considered high-confidence if they had been constructed using evidence from either PASA inputs or from ≥2 other prediction methods.

The transcriptome support shown in Supplementary Table 11 was calculated for each *P. glacialis* isolate by searching the high-confidence EvidenceModeler genes against a database of all RNA-Seq and IsoSeq transcripts (from the same isolate) using BLASTN. Genes were considered to have transcriptome support if they had a hit with >90% identity that covered >50% of the gene.

### Functional annotation of predicted genes

Protein domains were searched using pfam_scan.pl (v1.6; Pfam database release 31) at *E*-value < 0.001 following earlier studies^15, 41, 67^. Where required, proteins were queried using BLASTP against SwissProt and TrEMBL databases (UniProt release 2018_02) independently. Only the hits from the top 20 target sequences from each search were retained if *E* ≤ 10^-5^. Gene Ontology (GO; http://geneontology.org/) terms were assigned using the UniProt-GOA mapping (release 2019_05_07) database and the best UniProt hit with associated GO terms for each sequence.

Pfam domains in *P. glacialis* was assessed for enrichment against a background set using Fisher’s exact test, with correction for multiple testing using the Benjamini and Hochberg method^77^; a *p*-value ≤ 0.05 is considered significant. GO enrichment was conducted using the topGO R (v2.34.0)^78^ package, applying the Fisher’s Exact test with the ‘elimination’ methods to correct for the hierarchical structure of GO terms. The background used consisted of the available Symbiodiniaceae genomes (*Symbiodinium microadriaticum*^13^, *Breviolum minutum*^14^, *Cladocopium goreaui*^12^, *Fugacium kawagutii*^12^, *Symbiodinium tridacnidorum*^15^ and *Cladocopium* sp. C92^15^) with the revised gene predictions from Chen et al.^37^.

### Analysis of completeness of assembled genomes and predicted proteins

Completeness of the predicted genes in *P. glacialis* was assessed using BUSCO v3.1.0 (--mode proteins)^79^ with the alveolate_stramenophiles_ensembl, Eukaryota_odb9 and protists_ensembl datasets (retrieved 22 September 2017), BLASTP searches (*E* ≤ 10^-5^) using the same three BUSCO datasets and BLASTP searches *(E* ≤ 10^-5^) using the protein orthologs from the Core Eukaryotic Genes dataset^24^ (Supplementary Table 4).

Completeness of the assembled genomes of *P. glacialis* was assessed using BUSCO v3.1.0 (--mode proteins)^79^ and TBLASTN searches (*E ≤* 10^-5^) using the same three BUSCO datasets and TBLASTN searches (*E ≤* 10^-5^) using the protein orthologs from the Core Eukaryotic Genes dataset^24^ (Supplementary Table 4). The modified version of Augustus used for gene prediction was used for the BUSCO analysis as well.

### Identification of *P. glacialis* sequences in the TARA database

The Ocean Microbial Reference Gene Catalogue was retrieved from ftp://ftp.sra.ebi.ac.uk/vol1/ERA412/ERA412970/tab/OM-RGC_seq.release.tsv.gz. Genes classified as being from “Dinophyceae” or that were from the kingdom “undef” were extracted and searched against the genome of both *P. glacialis* isolates using BLASTN (at default parameters). Genes (i.e. the query) were retained if they had a hit in the genome sequences, in which the aligned region covered >75% of the query length at >95% identity. The retained genes were then searched against the non-redundant (nr) nucleotide database at NCBI for further verification of their origins (20 May 2019). No *P. glacialis* sequences were identified this way.

### Comparison of predicted proteins and genome-sequence similarity between *P. glacialis* isolates

Comparison between the protein sequences of CCMP1383 and CCMP2088 was conducted using BLASTP (*E* ≤ 10^-5^; Fig. 3a). For each isolate, protein sequences that do not share similarity to those of the counterpart isolate were identified. For these proteins, the corresponding coding gene sequences were searched (BLASTN) against the transcripts of the counterpart isolate; we consider a shared sequence similarity of >90% identity covering >50% of the query as significant. Tandem repeated genes were identified with MCScanX^80^ (intra-species; -b 1) using hits from BLASTP (*E* ≤ 10^-10^) with query or subject coverage >50%.

Sequence similarity between the genomes of the *P. glacialis* isolates was assessed using non-repeat regions of the genome. Repeat features predicted using RepeatModeler and RepeatMasker were excluded from the analysis; regions between repeats that were ≤10 bp of each other were also removed. From the remaining non-repetitive regions, only those ≥100bp and with ≤10 ambiguous (“N”) bases were used as query in a BLASTN (-dust no, *E* ≤ 10^-10^) search against the genome of the other isolate. The top hit of each sequence was retained for further analysis.

### Inference of homologous protein sets among Suessiales

Putatively homologous protein sets were constructed using OrthoFinder v2.3.3 (inflation 1.5)^81^ with sequence similarity computed with DIAMOND v0.9.24^82^. We defined dark homologous protein sets using the same criteria as Stephens et al. ^41^ but excluding hits from any sequences with functions described as “Uncharacterized”.

### Functional classification of rhodopsin

Predicted proteins of *P. glacialis* with top hits described as “Rhodopsin” in UniProt were retrieved. The specific type for each identified *P. glacialis* rhodopsin was identified using a BLASTP (*E ≤* 10^-5^) search against the known proton-pump (ABV22426, ADY17811) and sensory type (ADY17810, KF651052, KF651053, KF651054, KF651055) sequences from *Oxyrrhis marina.* The top hit for each query sequence was used to assign type.

### Phylogenetic inference of DUF3494 domains

A comprehensive set of DUF3494 domain-encoding proteins was collected from the transcriptomes of *Heterocapsa arctica* CCMP445, *Peridinium aciculiferum* PAER_2, *Scrippsiella hangoei* like-SHHI_4 and *Scrippsiella hangoei* SHTV5 (retrieved from Microbial Eukaryote Transcriptome Sequencing Project (MMETSP)^83^) for comparison against those predicted in both *P. glacialis* isolates. Predicted DUF3494-encoding proteins from the bacteria associated with *P. glacialis* were found with pfam_scan.pl (see above). DUF3494 domain regions were extracted from the proteins if they covered >50% the length of the Pfam DUF3494 domain HMM. DUF3494 domains from the Pfam_Full dataset (retrieved 14 April 2019) were retrieved. Identical sequences within each dataset were removed using cd-hit (-c 1.00 -n 5)^71^. All DUF3494 domains and domain regions were aligned using MAFFT v7.407 (--localpair --maxiterate 1000)^84^, from which a Maximum Likelihood tree was constructed using IQ-TREE v1.6.10 (-m MFP -msub nuclear -bb 2000 - nm 2000)^53, 85, 86^. Support of nodes in the inferred tree was determined using 2000 ultrafast bootstraps^53^.

### Analysis of simple repeats and multi-copy genes

The *de novo* repeat families identified by RepeatModeler during gene prediction were scrutinised for the presence of multi-copy genes. Unclassified repeat families (type unknown) were compared using BLASTN (*E ≤* 10^-5^) against the final gene models. Queries (repeat families) with >80% of their sequence, or the sequence of the subject (predicted genes), being covered in the BLAST hit were retained. This strategy (considering cover of both query and subject) is designed to capture cases where either the whole repeat is contained within a gene (repetitive exon) or a whole gene in contained within a larger repeat.

To specifically assess the presence of simple repeats in the assembled genomes, RepeatMasker was re-run on each genome using the default library (RepeatMasker database release 20170127) searching for just simple repeats (-noint). Repeats of type (TTG)n, (TGT)n, (GTT)n, (AAC)n, (ACA)n, and (CAA)n are all derived from the same pattern and thus are considered interchangeable for the purposes of this study. Overlapping repeats of these types were merged and their total length was reported as the coverage of the TTG repeat. 3-mers were extracted from the cleaned genome assembly using kmercountexact.sh from the bbmaps tool suite (Supplementary Table 5). The quality trimmed and merged genome reads were sampled at 5% before 3-mers were extracted (reformat.sh samplerate=0.05, 3-mers extracted using kmercountexact.sh). This was done to prevent 3-mer counts from exceeding the maximum value a 32 bit integer can store.

### Analysis of spliced leader sequences

Polished PacBio IsoSeq sequences that contained the dinoflagellate spliced leader sequence (CCGTAGCCATTTTGGCTCAAG) were identified using BLASTN (-max_target_seqs 1000000 -task blastn-short -evalue 1000). Only sequences with hits that start ≤5 bp from their 5′-end, ended ≥20 bp along the DinoSL sequence, had zero gap openings and a maximum of one mismatch were considered to contain the spliced leader sequence. Relic DinoSL sequences were identified by BLASTN (-max_target_seqs 1000000 -task blastn-short -evalue 1000), using the full DinoSL and a relic sequence joined together as the query^30^. Multiple relic DinoSL were identified using the full DinoSL and multiple relic DinoSL sequences joined together. Sequences were considered to contain a relic DinoSL if they had a hit that started within the first 11 bases of the query sequence (allows for truncation of the transcript), within the first 5 bases of the transcript, and finished within 5 bases of the end of the query sequence.

## Supporting information

Supplementary Tables 1 through 22

## Acknowledgements

T.G.S. was supported by an Australian Government Research Training Program (RTP) Scholarship. D.W.B and Y.C. are supported by a Human Frontier Science Program grant (RGP0030). This project was supported by two Australian Research Council grants (DP150101875 awarded to M.A.R., C.X.C. and D.B., and DP190102474 awarded to C.X.C. and D.B.), and the computational resources of the National Computational Infrastructure (NCI) National Facility systems through the NCI Merit Allocation Scheme (Project d85) awarded to C.X.C. and M.A.R.

## Author contributions

T.G.S., M.A.R., and C.X.C. conceived the study; T.G.S., R.A.G.P., C.X.C., M.A.R., D.B. and D.W.B. designed the analyses and interpreted the results; C.X.C. maintained the dinoflagellate cultures; C.X.C. and A.R.M. extracted biological materials for sequencing; Y.C. generated the long-read libraries for genome and full-length transcriptome sequencing; T.G.S. conducted all computational analyses, prepared all figures and tables, and prepared the first draft of the manuscript; all authors prepared, wrote, reviewed, commented on and approved the final manuscript.

## Data availability

All sequence data generated from this study are available at the NCBI Short Read Archive (SRA) BioProject accession PRJEB33539, with SRA accessions ERS3790104 and ERS3790106 for CCMP1383, and ERS3790105 and ERS3790107 for CCMP2088. The assembled genomes, predicted gene models and proteins from both *P. glacialis* isolates are available at: https://cloudstor.aarnet.edu.au/plus/s/Nx08JEMt7FjK3zY.

## Supplementary Information

### Supplementary Note

**Supplementary Fig. 1:** GenomeScope 21-mer profile for CCMP2088.

**Supplementary Fig. 2:** Interspersed repeat landscape and proportion of distinct repeat classes in the assembled genome of CCMP2088, relative to sequence divergence in Kimura substitution level.

**Supplementary Fig. 3:** Relationship between length of intergenic regions and their coverage by repeats for the predicted genes from (A) CCMP1383 and (B) CCMP2088. The red trend line was constructed using a moving average with a window size of 250.

**Supplementary Fig. 4:** An example of a genome region containing genes nested within the long introns of a putative alanine–tRNA ligase (from scaffold CCMP1383_scf7180000588947). The EvidenceModeler predicted genes, mapped IsoSeq transcripts and mapped RNA-Seq transcripts are shown in the green, red and blue boxes.

**Supplementary Fig. 5:** Conserved synteny between the two sequenced bacterial scaffolds and the published (A) *Paraglaciecola psychrophila* strain 170T (GenBank NC_020514) and (B) *Sphingorhabdus* sp. YGSM121 (GenBank NZ_CP022548) genomes. Syntenic regions between the two sequences are shown with ribbons; red representing direct and green represents inverted regions.

Supplementary Tables 1 through 22

Supplementary Data 1

