## Supplementary Note for "*Polarella glacialis* genomes encode tandem repeats of single-exon genes with functions critical to adaptation of dinoflagellates"

**Supplementary Note**
**Supplementary Figures 1 through 5**
**Supplementary Tables 1 through 22**
**Supplementary Data 1**

### Supplementary Note

#### *Bacterial species associated with P. glacialis in the environment*

Two bacterial genomes were generated as part of our effort in sequencing the genomes of *P. glacialis*. A CCMP1383 scaffold (5,892,869 bp) from the preliminary (short-read-only) assembly mapped at 93.8% sequence identity to the genome of *Paraglaciecola psychrophila* strain 170T (GenBank NC_020514, 5,413,691 bp; gamma-Proteobacteria), and another (3,839,769 bp) at 83.9% sequence identity to the genome of *Sphingorhabdus* sp. YGSM121 (GenBank NZ_CP022548, 3,864,176 bp; alpha-Proteobacteria). These two scaffolds show strong conserved synteny to the published genomes, with a few structural rearrangements (Supplementary Figure S5). *P. psychrophila* 170 was isolated from the Arctic[1], and *Sphingorhabdus* sp. YGSM121 from temperate sea sediment near South Korea (GenBank NZ_CP022548). Similarly, the three largest contaminant scaffolds (total 4,274,053 bp) in the preliminary CCMP2088 assembly have likely originated from the Arctic *Maribacter arcticus* (GenBank GCF_900167935.1, 4,211,145 bp; Bacteroidetes/Chlorobi) isolate. Therefore, these bacterial species are likely microbial associates of *P. glacialis* in the origin environment (i.e. part of the *P. glacialis* holobiont) where the two isolates were first isolated, but this notion remains to be investigated.

#### *Mitochondrial genomes*

In the assembled genomes of CCMP1383 and CCMP2088, 696 and 552 sequences, respectively were identified as being putative mitochondrial, totalling over 20 Mbp of sequence in each isolate (Supplementary Table S20). The longest mitochondrial sequence (827,212 bp), recovered from CCMP2088, putatively encodes 77 genes. This sequence is longer and does not conform to the same mitochondrial gene structure previously reported in dinoflagellates[2-4]. Based on BLASTN search ($E \leq 10^{-5}$) of each putative mitochondrial sequence within an isolate against those of the same isolate (excluding self-hits), >99.7% by length of these sequences are redundant; this percentage is 12.5% in CCMP1383 (and 50.2% in CCMP2088) when considering only hits covering >10 Kbp. This result suggests that these sequences within each isolate share little contiguity, and that they may undergo (or have undergone) frequent structural rearrangements.

By length, 86.6% of the putative mitochondrial sequences in CCMP1383 share significant similarity to 91.1% of their counterpart in CCMP2088, all of these aligned regions are <2

Kbp. This results suggests that the structural rearrangement we observed in each isolate is independent (and potentially active), resulting in significant structural differences between the two isolates.

The 3′ and 5′ ends of mitochondrial encoded genes in both isolates appear to be poorly conserved compared to the available "complete" mitochondrial genes of *Breviolum minutum* (extracted from GenBank accessions LC002801.1 and LC002802.1). For instance, the *cox1* and *cox3* genes in *P. glacialis* are truncated by ~30 and ~40 bp, respectively at the 5′-end; the *cox*3 gene is truncated by ~10-70 bp at the 3′-end.

The sequence similarity shared among mitochondrial genomes of dinoflagellates was further assessed using the two *B. minutum* mitochondrial genome sequences[4] in a BLASTN search ($E \leq 10^{-5}$) against putative mitochondrial sequences from CCMP1383, CCMP2088, *C. goreaui* and *F. kawagutii*[2]. Only eight sequence regions (total length 3,682 bp) of the *B. minutum* mitochondrial genomes were found to be conserved across all four other isolates (Supplementary Table S21). Three of these regions were identified as the mitochondrial encoded genes, two regions as pseudogenes, two regions represent fragments of the large subunit rRNA, and one region is uncharacterised. The function of the conserved non-genic regions remains to be further explored.
