## Supplementary figures and images for "*Polarella glacialis* genomes encode tandem repeats of single-exon genes with functions critical to adaptation of dinoflagellates"

### Supplementary Fig. 1

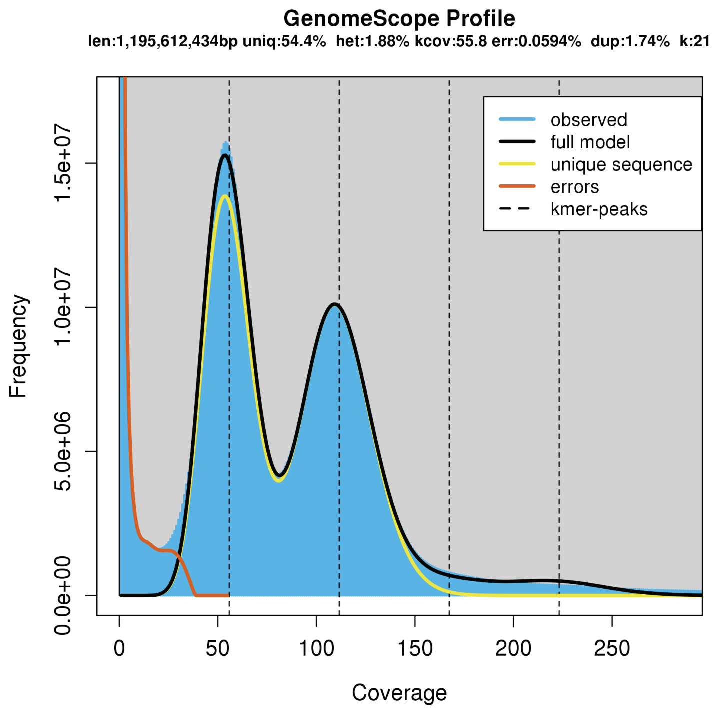


**Supplementary Figure 1.** GenomeScope 21-mer profile for CCMP2088.
