## Supplementary Fig. 2 for "*Polarella glacialis* genomes encode tandem repeats of single-exon genes with functions critical to adaptation of dinoflagellates"

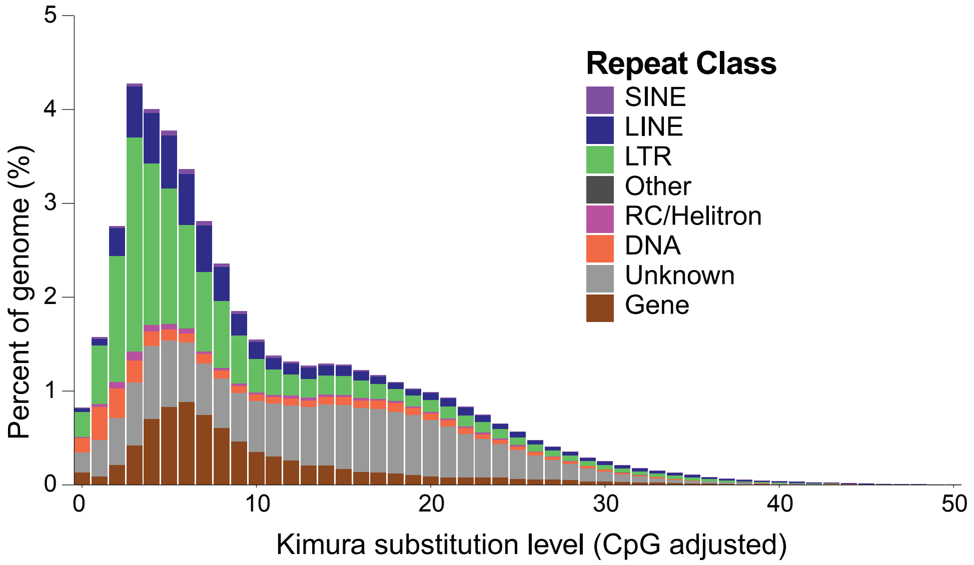


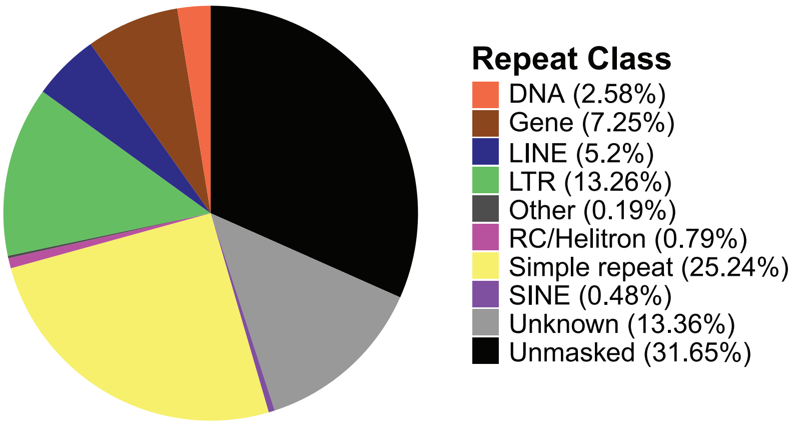


**Supplementary Figure 2.** Interspersed repeat landscape and proportion of distinct repeat classes in the assembled genome of CCMP2088, relative to sequence divergence in Kimura substitution level.
