## Supplementary Fig. 3 for "*Polarella glacialis* genomes encode tandem repeats of single-exon genes with functions critical to adaptation of dinoflagellates"

**a**

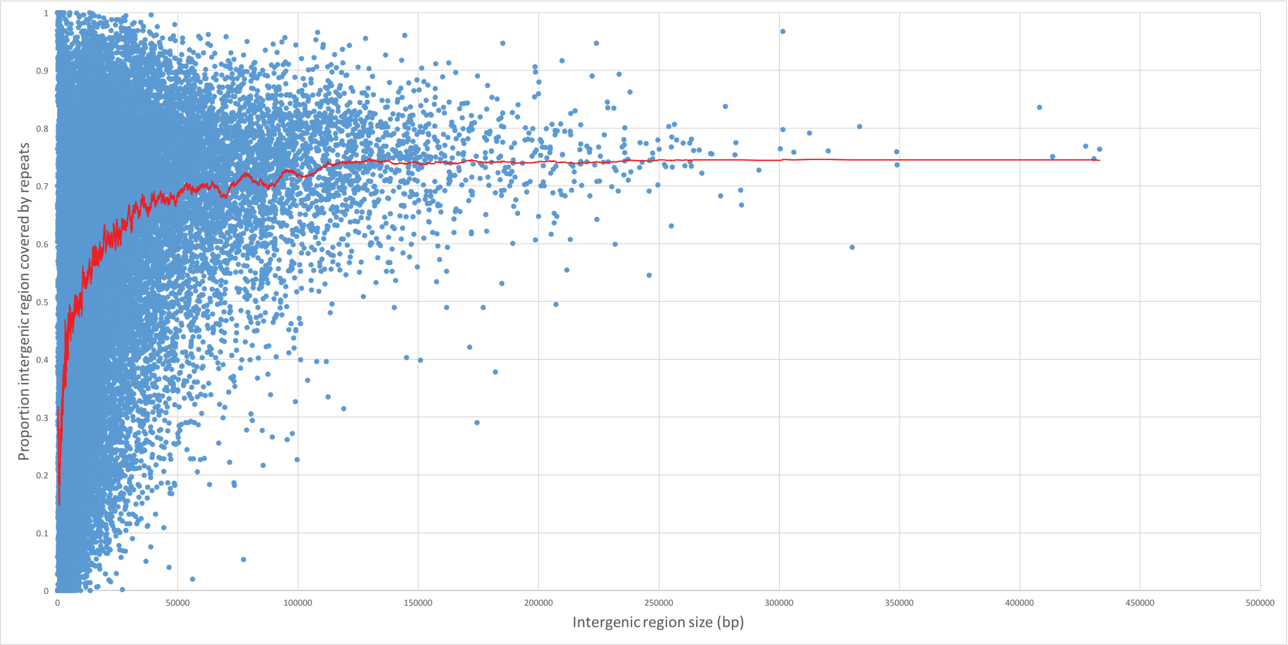


**b**

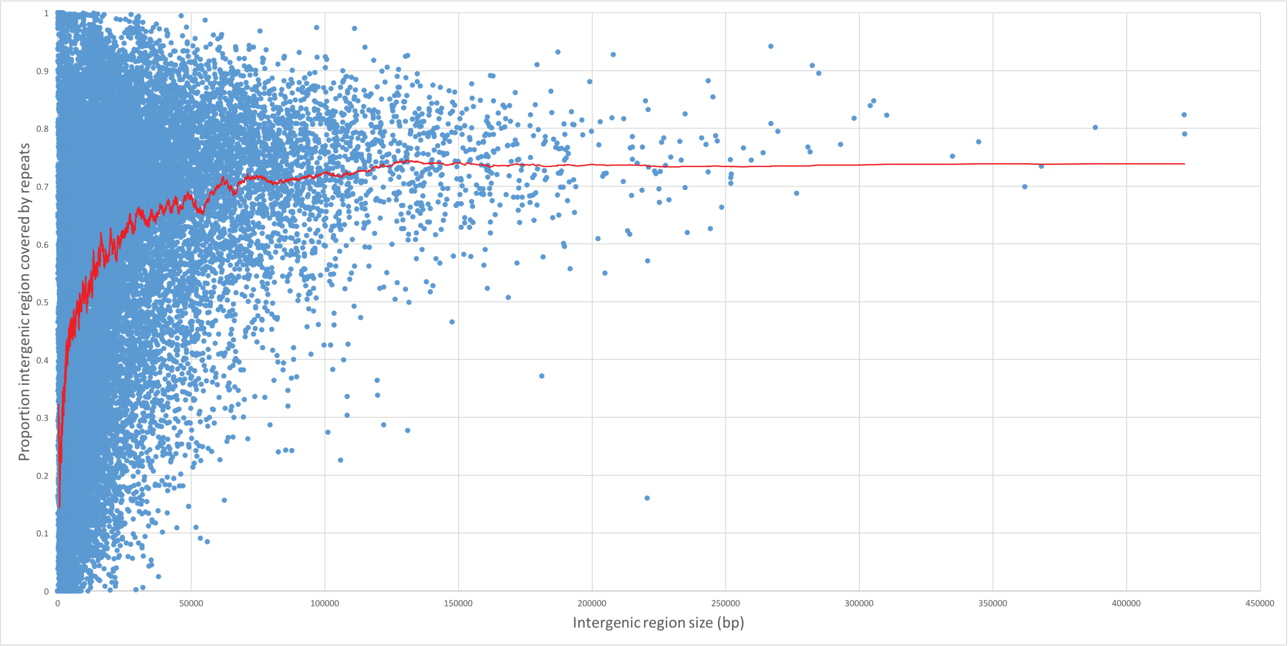


**Supplementary Figure 3.** Relationship between length of intergenic regions and their coverage by repeats for the predicted genes from (a) CCMP1383 and (b) CCMP2088. The red trend line was constructed using a moving average with a window size of 250.
