## Supplementary Fig. 4 for "*Polarella glacialis* genomes encode tandem repeats of single-exon genes with functions critical to adaptation of dinoflagellates"

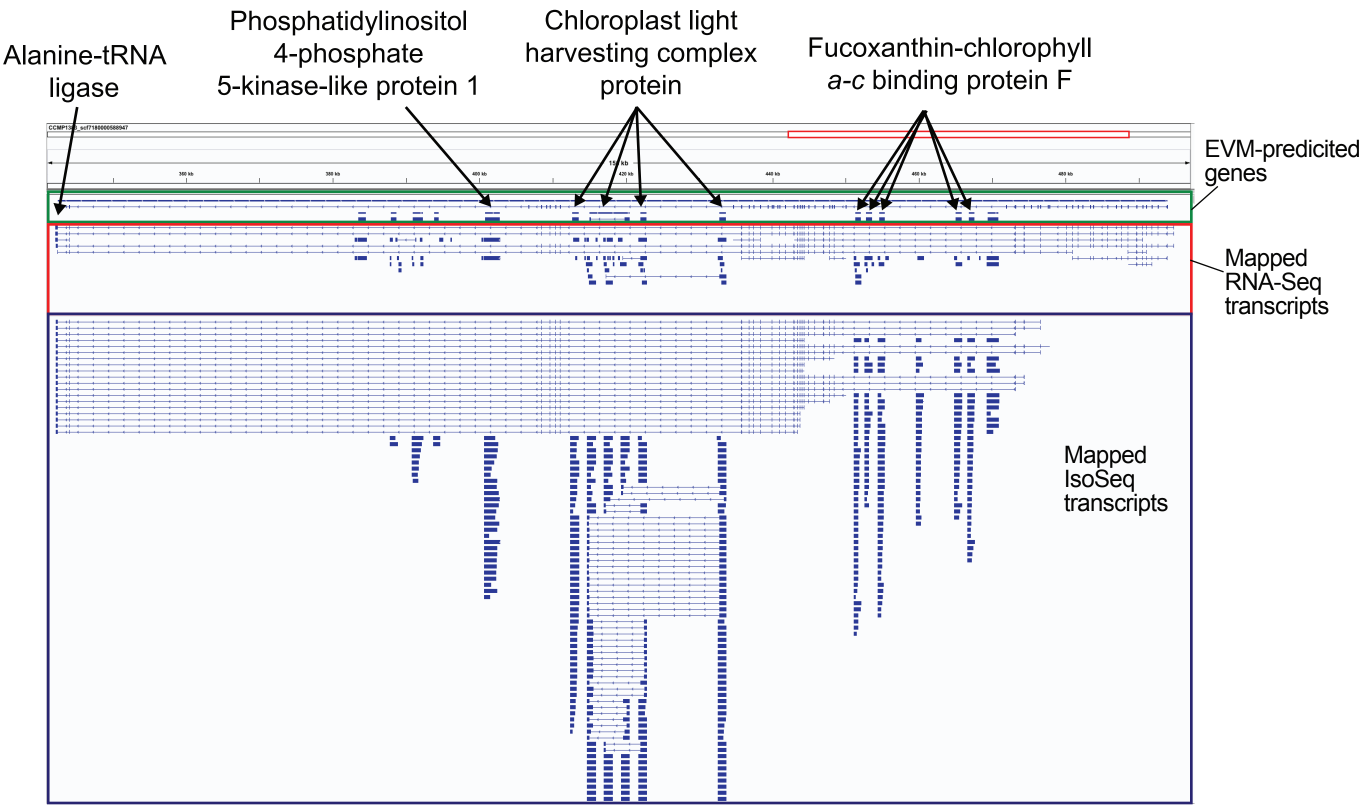


**Supplementary Figure 4.** An example of a genome region containing genes nested within the long introns of a putative alanine-tRNA ligase (from scaffold CCMP1383_scf7180000588947). The EvidenceModeler predicted genes, mapped IsoSeq transcripts and mapped RNA-Seq transcripts are shown in the green, red and blue boxes.
