## Supplementary Fig. 5 for "*Polarella glacialis* genomes encode tandem repeats of single-exon genes with functions critical to adaptation of dinoflagellates"

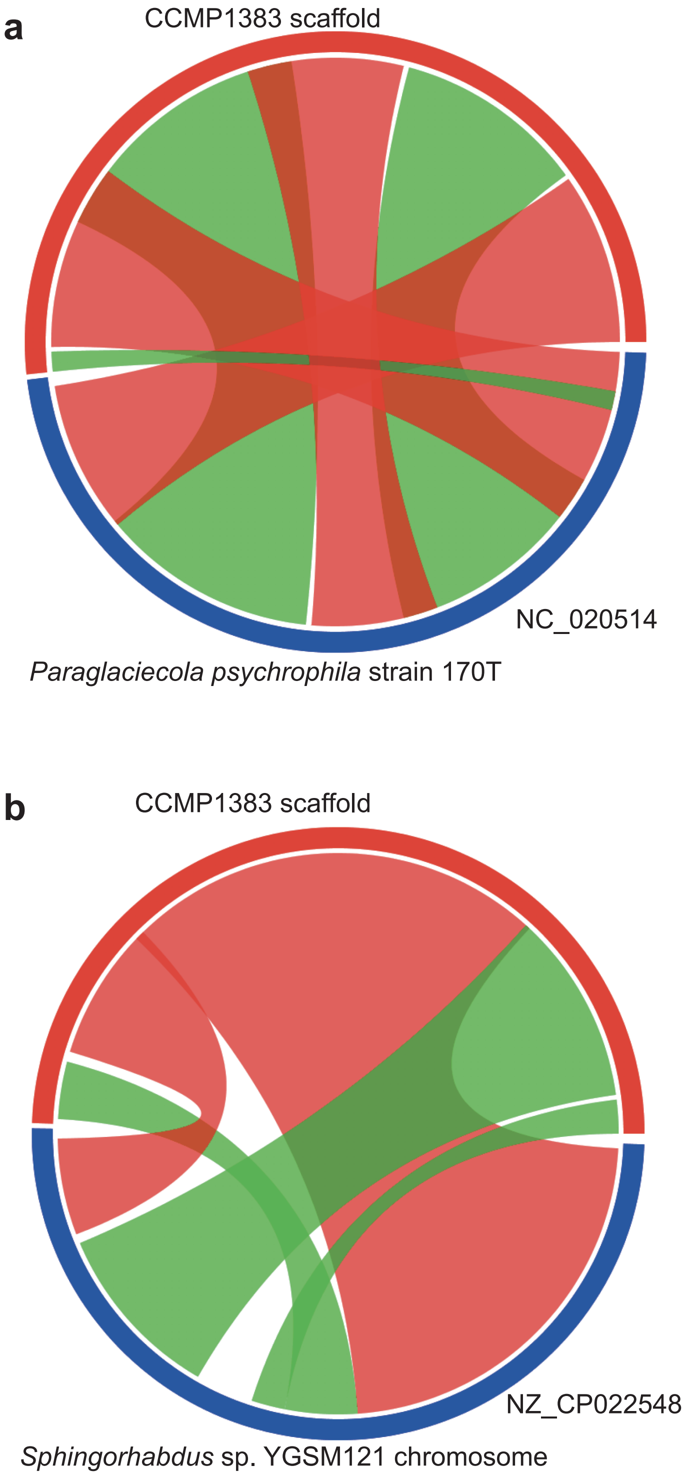


**Supplementary Figure 5.** Conserved synteny between the two sequenced bacterial scaffolds and the published (a) *Paraglaciecola psychrophila* strain 170T (GenBank NC_020514) and (b) *Sphingorhabdus* sp. YGSM121 (GenBank NZ_CP022548) genomes. Syntenic regions between the two sequences are shown with ribbons; red representing direct and green represents inverted regions.
